# DeepMed: A unified, modular pipeline for end-to-end deep learning in computational pathology

**DOI:** 10.1101/2021.12.19.473344

**Authors:** Marko van Treeck, Didem Cifci, Narmin Ghaffari Laleh, Oliver Lester Saldanha, Chiara M. L. Loeffler, Katherine J. Hewitt, Hannah Sophie Muti, Amelie Echle, Tobias Seibel, Tobias Paul Seraphin, Christian Trautwein, Sebastian Foersch, Tom Luedde, Daniel Truhn, Jakob Nikolas Kather

## Abstract

The interpretation of digitized histopathology images has been transformed thanks to artificial intelligence (AI). End-to-end AI algorithms can infer high-level features directly from raw image data, extending the capabilities of human experts. In particular, AI can predict tumor subtypes, genetic mutations and gene expression directly from hematoxylin and eosin (H&E) stained pathology slides. However, existing end-to-end AI workflows are poorly standardized and not easily adaptable to new tasks. Here, we introduce DeepMed, a Python library for predicting any high-level attribute directly from histopathological whole slide images alone, or from images coupled with additional meta-data (https://github.com/KatherLab/deepmed). Unlike earlier computational pipelines, DeepMed is highly developer-friendly: its structure is modular and separates preprocessing, training, deployment, statistics, and visualization in such a way that any one of these processes can be altered without affecting the others. Also, DeepMed scales easily from local use on laptop computers to multi-GPU clusters in cloud computing services and therefore can be used for teaching, prototyping and for large-scale applications. Finally, DeepMed is user-friendly and allows researchers to easily test multiple hypotheses in a single dataset (via cross-validation) or in multiple datasets (via external validation). Here, we demonstrate and document DeepMed’s abilities to predict molecular alterations, histopathological subtypes and molecular features from routine histopathology images, using a large benchmark dataset which we release publicly. In summary, DeepMed is a fully integrated and broadly applicable end-to-end AI pipeline for the biomedical research community.

## Introduction

### End-to-End Deep Learning in Computational Pathology

Histopathology slides stained with hematoxylin and eosin (H&E) are ubiquitously available for virtually every single patient with a solid tumor.^1^ H&E tissue slides are indispensable for making a disease diagnosis. Beyond that, they are broadly used to derive qualitative and quantitative biomarkers for translational and basic cancer research studies.^2^ Artificial intelligence (AI), specifically Deep Learning (DL) with convolutional neural networks (CNNs) can be used to automatically analyze digitized whole slide images (WSIs) of H&E slides and can yield quantifiable information beyond the capabilities of human experts. In the last four years, multiple research groups have shown that DL methods can predict high-level concepts such as the presence of specific genetic mutations^3,4^, gene expression^5^, whole genome duplications^6^, patient survival^7^ and treatment response^8^ from H&E WSI. Since the first publication in 2018^9^ demonstrated a robust end-to-end workflow, more than one hundred academic studies have used similar approaches^2^. However, unlike in other areas of bioinformatics, there are currently no standard pipelines for end-to-end-DL in computational pathology. Therefore, virtually all research teams who are active in this field have implemented their own pipeline with highly similar setups. For example, multiple analysis pipelines have been developed between 2018 and 2021 to predict the mutations of oncogenic driver genes from H&E WSI.^3,5,9–12^ The overall design of these pipelines is largely identical: they load a WSI, tessellate it into tiles, perform data augmentation and/or normalization, train a CNN, deploy the network on tiles from test patients and use an aggregation function to pool the tile-level predictions on a patient level.^9^

### Limitations of Previous Deep Learning Pipelines in Computational Pathology

Why do researchers re-implement essentially identical pipelines instead of re-using source codes of previous publications? A key reason is that published pipelines are not modular. Individual components of these pipelines are highly interconnected and cannot be easily changed without disrupting the overall workflow (**Figure 1A**). For example, many methods have been designed to train CNNs on a training set and test them on a designated test set.^9^ Others have used stratified cross-validation^4,13^ or Monte-Carlo cross-validation^14^ on a patient-level. Moving from one experimental design to another requires a multitude of upstream and downstream changes, related to data preprocessing, statistical metrics, visualization and essentially any component of the pipeline. Also, using the pipeline for different types of input data (for example, prediction of continuous instead of categorical values) disrupts the whole workflow from data loading, training to visualization of the results. Finally, end-to-end DL pipelines are being run on different types of hardware ranging from laptop computers with a single graphics processing unit (GPU) over workstations, in-house servers to commercial cloud computing services (**Figure 1B**) with Windows or Linux operating systems. Current processing pipelines cannot be easily deployed on these different types of hardware and operating systems. We aimed to address these issues and developed DeepMed, a modular, extensible, versatile, easily usable, powerful DL pipeline for end-to-end computational pathology in translational and basic research.

**Figure 1:**
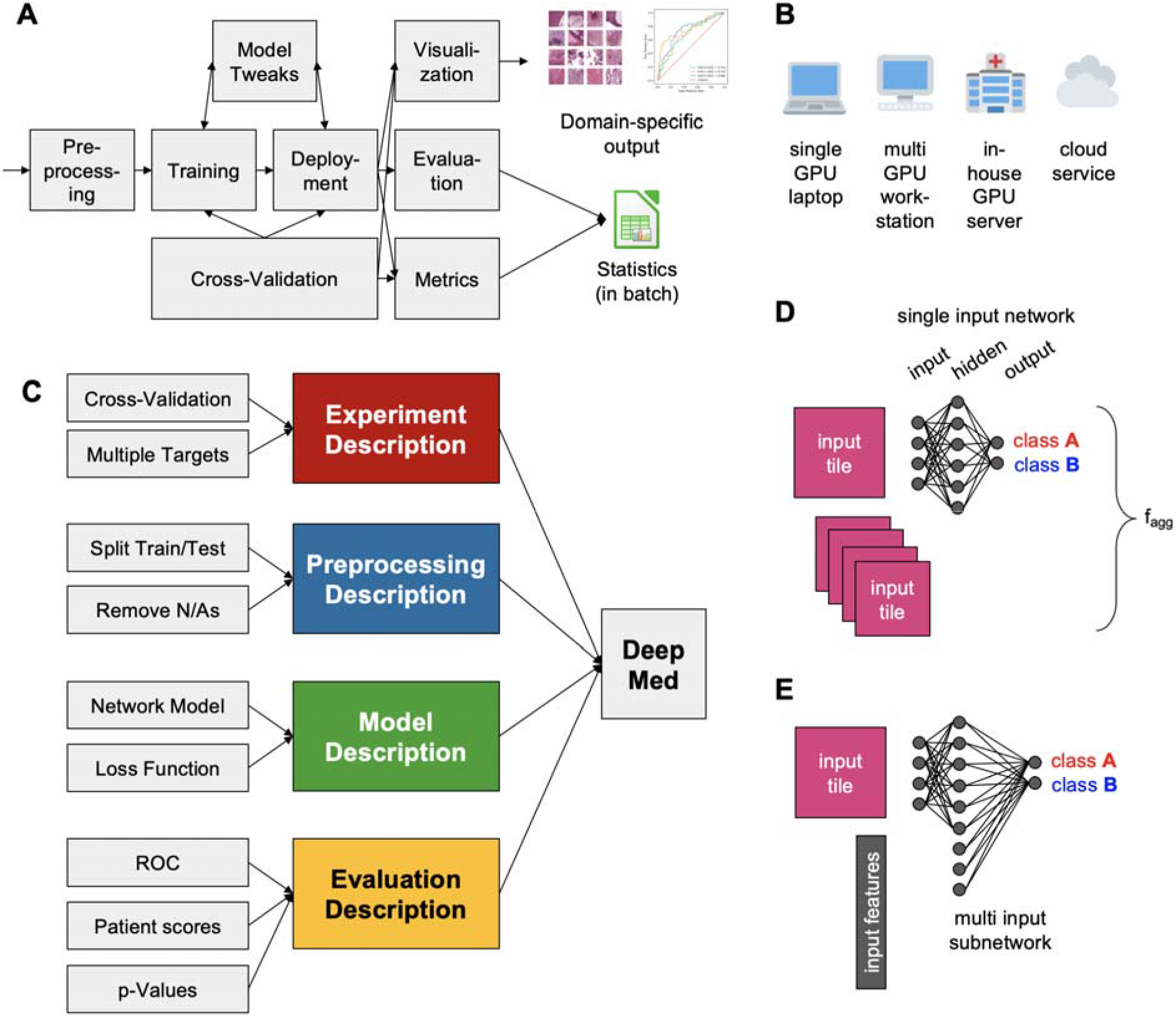
Overview of the DeepMed concepts. (**A**) The main components of a typical deep learning pipeline for biomedical image analysis, (**B**) The computer systems that run deep learnin**g** pipelines, (**C**) The modular structure of DeepMed that serves different functions to conduct an end-to-end bioimage analysis, (**D**) The network representation of a single model’s training on a bag of image tiles in the *simple run* mode, Abbreviations: f_agg = aggregation function. (**E**) The network representation of a single model’s training with multiple inputs on the *simple run* mode. Only a single tile and a single set of features is visually shown, although in reality, the model processes a bag of image tiles and a corresponding set of input features.

### Development, Application and Validation of the Protocol

DeepMed is a pipeline that integrates a multitude of algorithms which were developed and evaluated in end-to-end computational pathology. Unlike previously published pipelines, DeepMed includes all commonly used variants of data loading, network training, statistics and visualization in a fully modular way (**Figure 1C**). DeepMed can be used for a wide range of problems including: simple classification and regression tasks on histological image data only,^3,9,13,15–18^, prediction of survival markers ^19^ (**Figure 1D**) and inclusion of additional non-image in the training process in a multi-modal way^20^ **(Figure 1E**). DeepMed enables researchers to conveniently use established methods and test dozens of hypotheses in a single cohort or multiple patient cohorts with minimal data preprocessing. At the same time, it offers a high degree of flexibility for developers who can use the robust backbone of DeepMed to try out new methods without the overhead of re-implementing a full end-to-end DL pipeline. Computational pathology is a fast-evolving field attracting much attention worldwide, evidenced by the increasing numbers of publications.^2^ Researchers must be able to iterate new ideas rapidly and scalably in order to keep up with new advances. DeepMed enables computational researchers to quickly try out many different approaches on large datasets. The modular composition of this pipeline allows programmers to easily adapt our code to fit their workflows. Furthermore, as new technologies or approaches become available, sections can be updated without changing the user-facing parts of the pipeline. Here, we present a comprehensive overview of setting up, using, validating and extending DeepMed and provide two benchmark datasets which can be used for many common problems in end-to-end computational pathology.

### Overview of the Algorithm and Workflow

From a user’s perspective, DeepMed is a convenient implementation of previously described algorithms which have been used for several medical image processing tasks. First, the user prepares three types of data (**Figure 2A**): A set of whole slide images (WSIs) in an openslide compatible format (https://openslide.org/), a clinical (“CLINI”) table which assigns *patients* (each row is one patient) to *targets* (e.g. mutational status of gene X) and a SLIDE table which assigns WSIs to patients (each row is one WSI). This data format was previously defined and extensively described in the “The Aachen Protocol for Deep Learning Histopathology”.^21^ Subsequently, the user starts the tessellation and, optionally, color normalization^22^ of WSIs, yielding individual image tiles saved on the hard disk. For example, this can be achieved with QuPath^23^ or with our own Python-based open source tessellation and normalization tool which is available as a commandline version (**Figure 2B**, https://github.com/KatherLab/preProcessing) and as a graphical user interface (GUI, https://github.com/KatherLab/preProcessing/GUI). Finally, the user defines an experiment in a Python script (“.py file”), as will be explained below (**Figure 2C**) and runs this script from the command line. Alternatively to creating a Python script, users can use the experimental DeepMed GUI for Microsoft Windows, which enables running DeepMed without any direct use of Python scripts (https://github.com/KatherLab/deepmed-gui). Without any further interaction with the user, DeepMed will preprocess the data (**Figure 2D**): it will parse the CLINI table, removing patients with missing values (which can be defined by the user as “NA”, “N/A”, “NaN” or other placeholders). In addition, it can restrict the analysis on subgroups (patients with a particular feature as defined in the CLINI table), it will collect the image tiles, perform train/test splits and plan the computing jobs for distribution to multiple GPUs. Subsequently, a neural network model (e.g., a model pretrained on Imagenet^24^) is trained on the tiles to predict the targets and will be saved for future deployment and/or directly deployed on a test set with known or unknown status of the target (**Figure 2E**). During training, the deep learning neural network learns to map a set of inputs to a set of outputs provided by the training data. In more detail, training allows the network to optimize weights the model uses to provide accurate predictions, i.e. predictions that exhibit a high concordance with the ground-truth. Performance of the model is evaluated by calculating the objective (loss) function. The typical loss function for categorical targets (binary/Multi-class) is Cross -Entropy (Binary Cross-Entropy [BCE] / Cross-Entropy [CE]) and the common loss functions for continuous targets are Mean Square Error (MSE) or Mean Absolute Error (MAE). By modifying the final layer of the DL neural network (one output node for continuous targets and *N* output nodes for *N* class categorical targets) and selecting the best loss function for optimization, it is possible to train DL algorithms for a wide range of targets. Finally, and most importantly for the end user, a range of statistics and visualizations can be generated (**Figure 2F**), including highly predictive image tiles^6^, receiver operating characteristic curves (ROCs)^9^, whole-slide prediction heatmaps^3^ and additional tile-level and patient-level statistics including AUROC, F1-score, p-values and others (shown in **Suppl. Table 2**).

**Figure 2:**
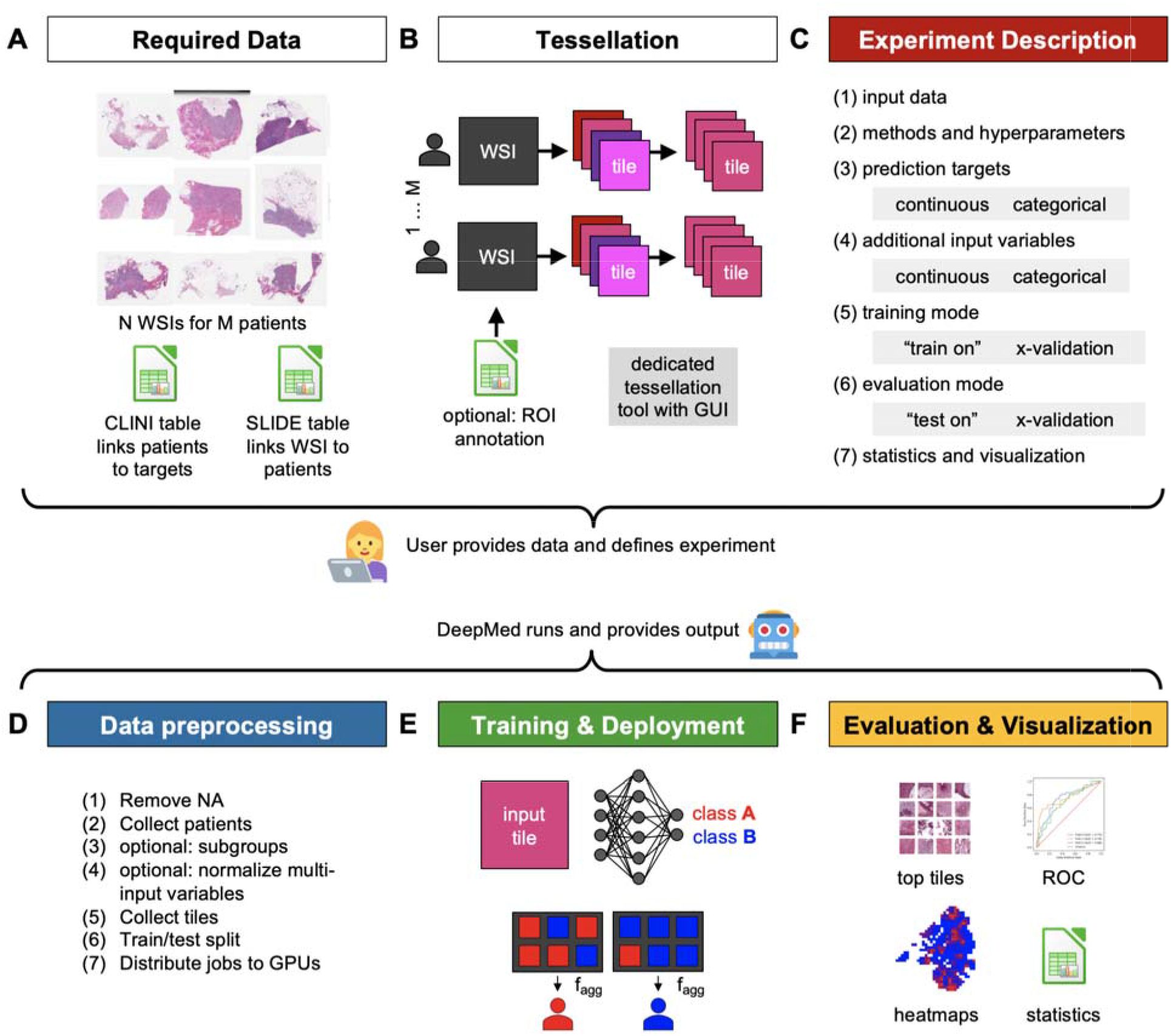
Overview of the DeepMed workflows from a user perspective. (**A**) The data to be curated prior to preprocessing, (**B**) The recommended preprocessing steps for WSIs which include tessellation and normalization, (**C**) The main parameters to be initialized in the experiment script or GUI of DeepMed, (**D**) The data preprocessing steps of DeepMed before the neural network training, (**E**) A rough framework of DeepMed that includes training deep learning mode**ls** with desired modes, evaluating them and pooling the statistics with an aggregation function on per-patient or -tile basis, (**F**) Example evaluation metrics provided by DeepMed. Abbreviations: WSIs: whole slide images, f_agg = aggregation function.

## Experimental Results

### Prediction of molecular features from breast cancer histology images

Here we present the results achieved for the main functionalities, summarized in **Table 1**, of DeepMed in benchmark datasets which are provided under an open access license at https://zenodo.org/record/5337009. As a benchmark task, we use prediction of pathological and molecular features in breast cancer based on slide-level (WSI-level) labels. This problem has been widely investigated in dedicated studies^8,11,25,26^ and as part of systematic pan-cancer studies^4–6^ and represents a clearly defined weakly supervised prediction task. We applied DeepMed to predict estrogen receptor (ER) status, histological subtype (ductal or lobular), TCGA gene expression subtype (Basal, LumA and LumB) and the density of tumor infiltrating lymphocytes (TILs).

**Table 1:** Key modules for all four branches of DeepMed.

| Experiment | Preprocessing | Model | Evaluation |
| --- | --- | --- | --- |
| 1. Within-cohort k-fold cross validation<br>2. Split a cohort into train and test cohort<br>3. Train on cohort A, deploy on cohort B<br>4. Parameterization allowing users to run consecutive experiments with different parameters | 1. Loading tiles and balancing classes by undersampling<br>2. Resize tiles to match network input size<br>3. Default transforms eg. data augmentations as rotation and flip | 1. Classification networks to predict categories<br>2. Regression networks to predict continuous values<br>3. Multi-input networks (integrate images and tabular data) | 1. AUROC, F1-Score and r<sup>2</sup><br>2. Receiver operating curve (ROC)<br>3. Heatmap visualization on whole slide level<br>4. Input tiles with the highest prediction score visualized as a montage<br>5. P-values for group comparisons |

We refer to the scripts to perform weakly supervised deep learning analysis with DeepMed analysis as “experiments” and give a detailed description of how to construct DeepMed experiments in the section “Materials, Methods and Procedure”. The first sample case experiment “train_and_deploy_multitarget.py” (**Full Script 4**) runs a DeepMed analysis that applies transfer learning with ResNet-18 on the Benchmark dataset, TCGA-BRCA-A2 (from Walter Reed National Military Medical Center, Bethesda, MD, USA). Subsequently, the resulting neural network model is deployed to make the predictions for three targets on the second, independent, test dataset, TCGA-BRCA-E2 (from Roswell Park Comprehensive Cancer Center, Buffalo, NY, USA). When we applied this workflow and evaluated the performance of the model on the test dataset, the classification performance was high, ranging from a patient-level AUROC of 0.656 for ER status to 0.860 for histological subtype (**Table 2**). All targets reached statistical significance (p<0.05).

**Table 2:** Aggregated statistics for multi-target deployment experiment.

| Target | Class | AUROC | Patient count | p value |
| --- | --- | --- | --- | --- |
| ER Status By IHC | Positive | 0.656 | 68 | 3.04E-02 |
| ER Status By IHC | Negative | 0.656 | 22 | 3.04E-02 |
| Histologic Type Name | Infiltrating Ductal | 0.860 | 74 | 8.00E-06 |
| Histologic Type Name | Infiltrating Lobular | 0.860 | 15 | 8.00E-06 |
| TCGA Subtype | BRCA.LumA | 0.889 | 47 | 0.00E+00 |
| TCGA Subtype | BRCA.LumB | 0.761 | 11 | 9.43E-04 |
| TCGA Subtype | BRCA.Basal | 0.839 | 15 | 3.00E-06 |
| TIL Regional Fraction | $[-\infty, 1.461)$, i.e. low | 0.777 | 51 | 3.00E-06 |
| TIL Regional Fraction | $[1.461, \infty)$, i.e. high | 0.777 | 36 | 3.00E-06 |

If an independent test dataset is not available, researchers can perform exploratory analyses with DeepMed using patient-level cross-validation. As a demonstration of this feature, we ran DeepMed analysis to evaluate the performance on unseen patients only on the training dataset TCGA-BRCA-A2, which is described in “crossvalidated_train_multitarget” (**Full Script 5**). In this within-cohort analysis, we found a moderate to high prediction performance for all targets, but statistical significance was only reached for the easiest task, prediction of histological subtype (p=0.023) (**Table 3**).

**Table 3:**
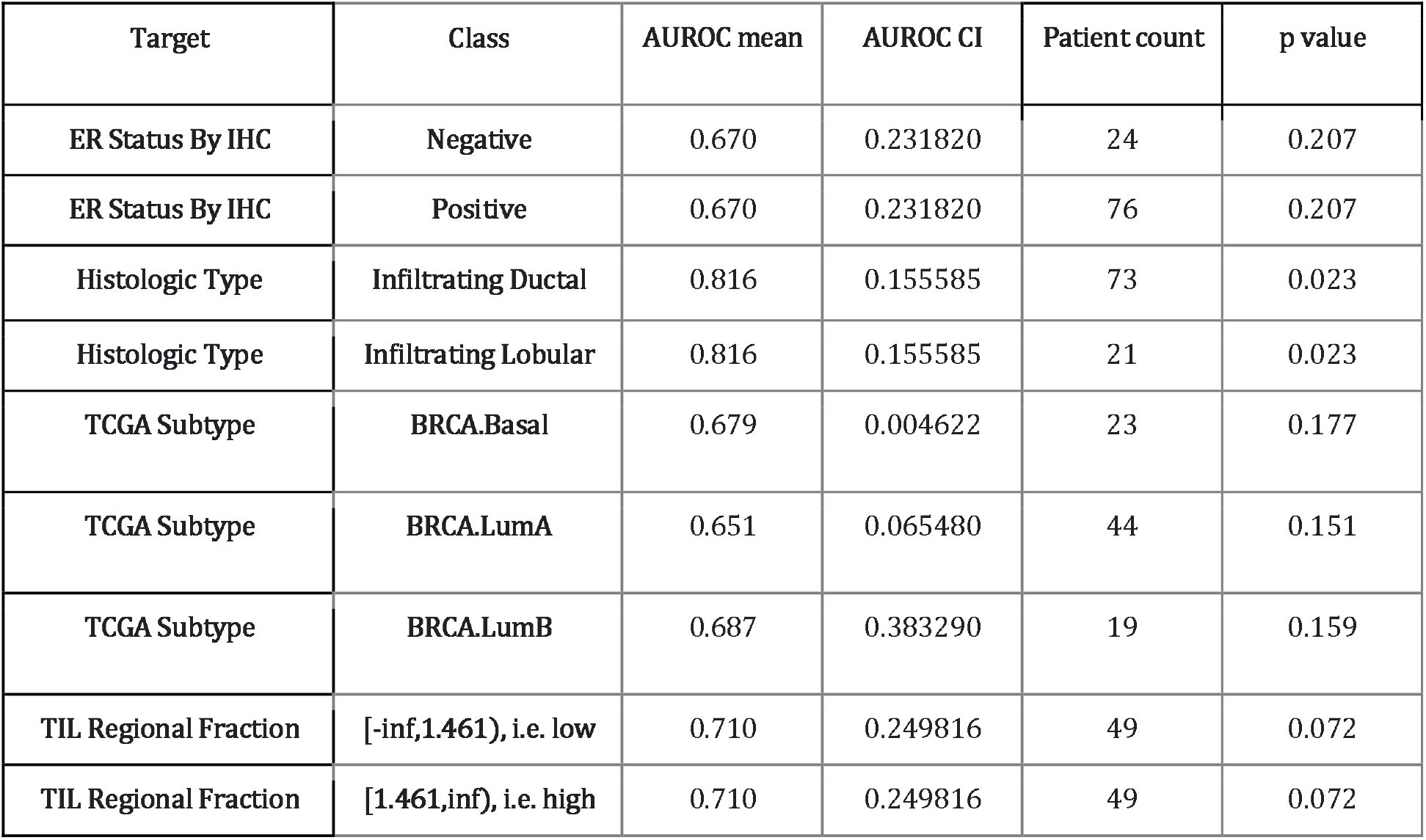
Aggregated statistics for multi-target 3-fold cross-validation experiment. CI: 95% confidence interval.

The receiver operating characteristic (ROC) curves that are among the output of the DeepMed analysis plotted by true positive rate (TPR) against the false positive rate (FPR) and demonstrating the performance of both cross-validated within-cohort and the train-and-deploy analyses are shown together in **Figure 4**. The ROC curves are created for each different class of predicted targets. In the case of cross-validated tasks, the curves for individual folds for each class are given in the same graph along with the calculated mean curve for the ease of interpretation of results. Furthermore, DeepMed can produce a single output image which includes a collage of tiles with highest prediction scores for each class (“top tiles”). In the top tiles-collages of classes, each row stores the tiles with the highest prediction scores from the patients with the highest prediction scores. Therefore, the size of the image is the requested number of patients times the number of tiles, which is 4 times 4 by default while these numbers can be changed in the experiment scripts. **Figure 5** shows the top tiles predicted by the models trained on the dataset TCGA-BRCA-A2 and deployed on the dataset TCGA-BRCA-E2. This type of visualization enables explainability of DL models, which is useful as a plausibility check as well as for discovery of new morphological biomarkers.^27^

**Figure 4:**
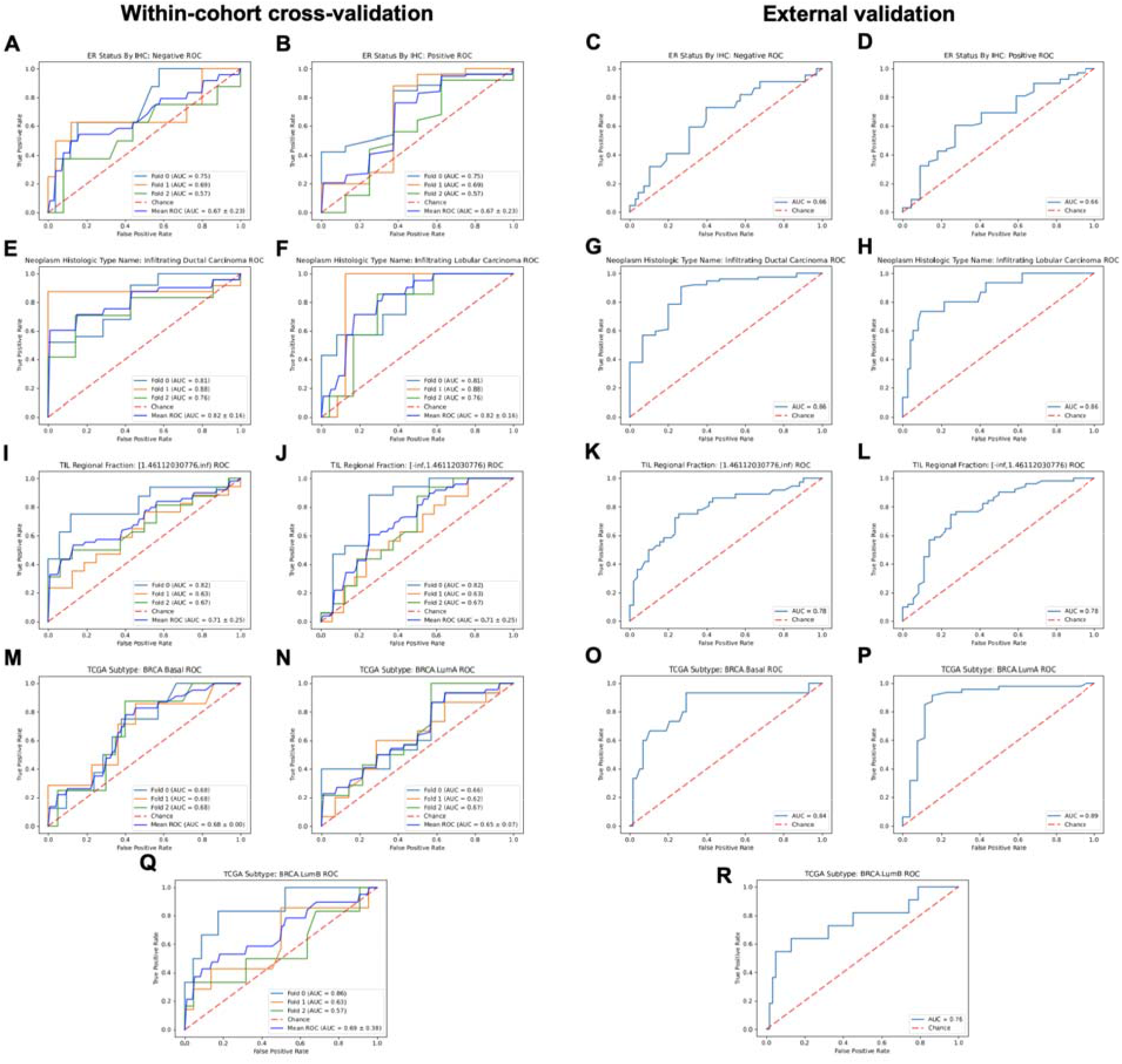
**The ROC curves of showing the classification performance of the model on** the negative class of ER Status **(A)** within cohort (TCGA-BRCA-A2) and **(C)** on the test dataset (TCGA-BRCA-E2), the positive class of the target ER status **(B)** within cohort and **(D)** on the test dataset; infiltrating ductal carcinoma class of histologic type name of neoplasm **(E)** within cohort and **(G)** on the test dataset; infiltrating lobular carcinoma class of histologic type name of neoplasm **(F)** within cohort and **(H)** on the test dataset; the class [1.461, inf] of the TIL regional fraction target (in this case, the range [1.461, inf] refers to the higher values in the continuous variable of TIL regional fraction, which was binarized at the median, 1.461) **(I)** within cohort and **(K)** on the test dataset; the class [-inf,1.461] of the TIL regional fraction target **(I)** within cohort and **(K)** on the test dataset; the TCGA subtype basal **(M)** within cohort and **(O)** on the test dataset; the TCGA subtype luminal A **(N)** within cohort and **(P)** on the test dataset; the TCGA subtype luminal B **(Q)** within cohort and **(R)** on the test dataset. Abbreviations: ER = estrogen receptor, TIL = tumor-infiltrating lymphocytes.

**Figure 5:**
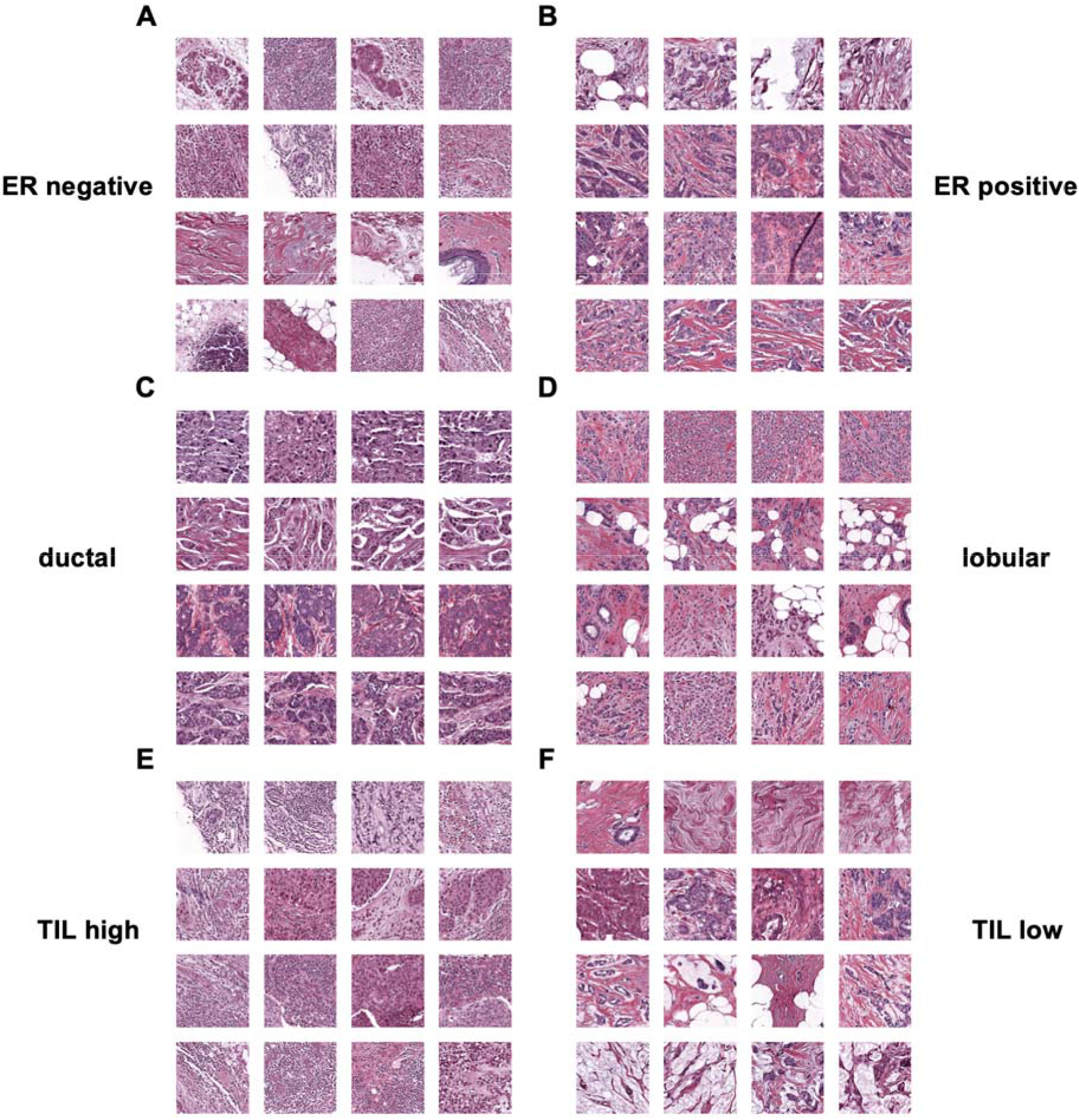
**The top predicted tiles of the top predicted patients by the models trained on TCGA-BRCA-A2 and deployed on TCGA-BRCA-E2 for (A)** the negative class and **(B)** the positive class of ER status; **(C)** infiltrating ductal carcinoma and **(D)** infiltrating lobular carcinoma of histologic type name of neoplasm; **(E)** high TILs (the class [1.461,inf]) and **(F)** low TILs (the class [-inf,1.461]) of the TIL regional fraction target (where the classes are artificially named upon discretizing them with the cutoff point 1.461); **(G)** the TCGA subtype basal, **(H)** the TCGA subtype luminal A **(I)** the TCGA subtype luminal B. These figures are generated by DeepMed with the default parameters: number of patients set to 4 and number of tiles set to 4, meaning that each row represents a patient that has a highest overall score while the tiles in the columns are the tiles with the highest scores from the respective patient.

Another strength of the DeepMed is the subgroup training functionality, which enables users to apply the DL analysis on a subset of patients of the original dataset. The subsets are defined by users in the experiment script.In “train_and_deploy_subgroup_based_TMB.py” (**Full Script 9**), we show how to train a model for prediction of ER status on subgroups of patients based on their tumor mutational burde (TMB, low and high as binarized at the median). The results show that ER status was predictable with an AUROC of 0.687 and 0.768 for TMB-high and TMB-low subgroups respectively with a p-value smaller than 0.05 in both (**Table 5**).

**Table 4:**
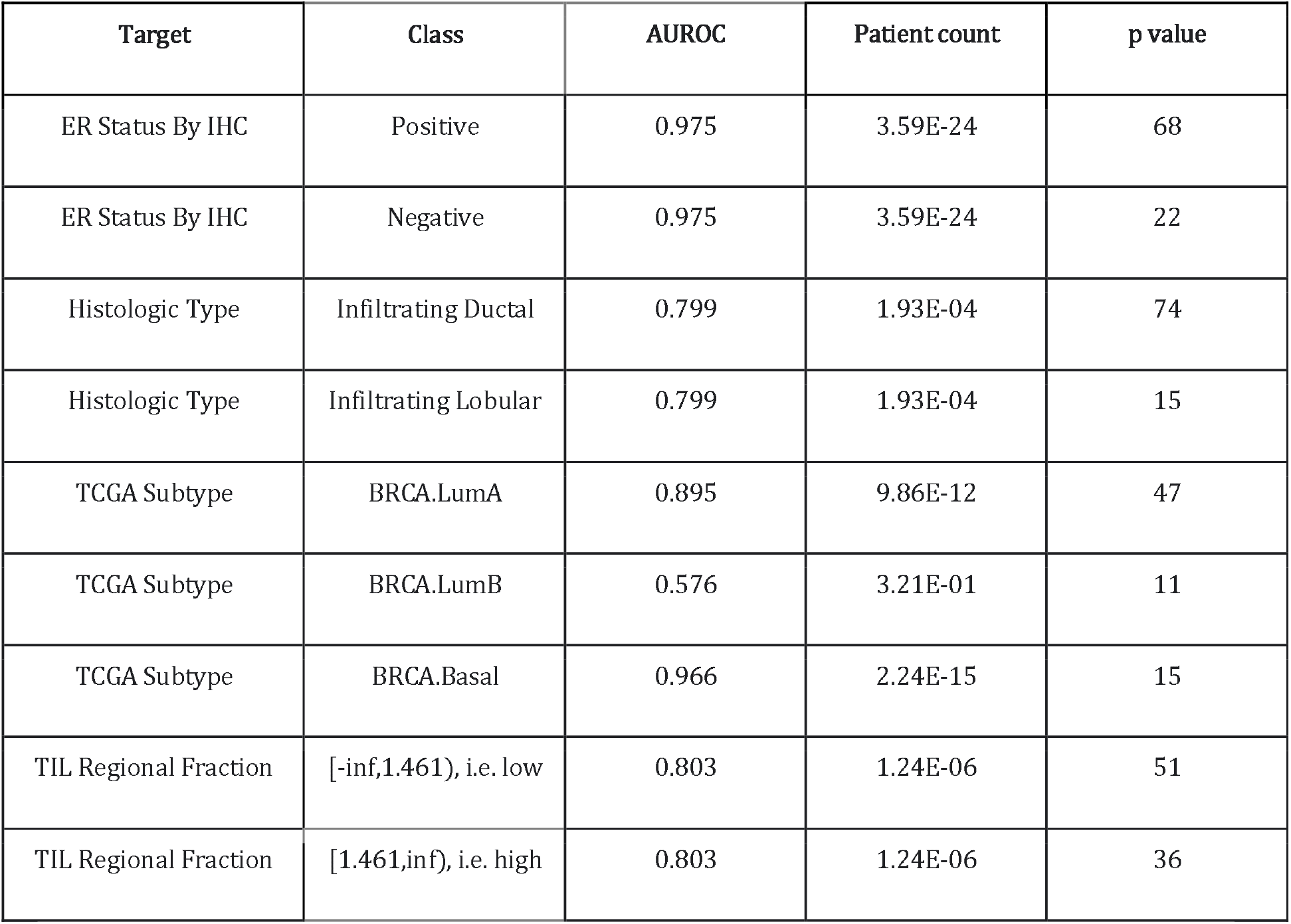
Aggregated statistics for multi-target deployment experiment experiment when HER2 and PR status are used as additional categorical inputs and the patient age is used as continuous input to the image data.

**Table 5:** Aggregated statistics deployment experiment targeting ER status in the subgroups with low and high tumor mutational burden (TMB) values respectively.

| Subgroup | Target | Class | AUROC | Patient count | p value |
| --- | --- | --- | --- | --- | --- |
| TMB high | ER Status By IHC | Positive | 0.687 | 31 | 2.97E-02 |
| TMB high | ER Status By IHC | Negative | 0.687 | 15 | 2.97E-02 |
| TMB low | ER Status By IHC | Positive | 0.765 | 34 | 4.30E-02 |
| TMB low | ER Status By IHC | Negative | 0.765 | 7 | 4.30E-02 |

These findings demonstrate the utility of DeepMed even in extremely small datasets of only 100 patients or less. Real-world applications of weakly supervised Deep Learning usually train and validate the models on thousands of cases^2,16,18,28,29^ and a continuous improvement of classifier performance has been demonstrated for higher patient numbers.^15,28^

### Prediction of molecular features from multi-modal data sources (multi-input mode)

In clinical decision making, healthcare providers rarely use only a single data type. Usually, different types of data, for example images and tabular data are used.^30^ DeepMed can integrate this multi-modal decision making by incorporating additional variables as an input to the neural network. We repeated the analyses of the breast cancer dataset and additionally provided the model with progesterone receptor (PR) status, HER2 status and age as input variables. The experiment script is shown in “train_and_deploy_multitarget_multiinput.py” (**Full Script 8**). We found that this addition of non-image information to the training markedly improves classifier performance in an external validation experiment (**Table 4**), showing that DeepMed can leverage tabular information to boost image-based prediction performance.

The parameterization mode of DeepMed has also been demonstrated with the multi-modality feature in “train_and_deploy_parameterizing.py” (**Full Script 10**). Parameterization mode provides users with the opportunity of running the same experiment with an unlimited number of different parameters separately and returns the results in the same project folder with an overall statistics report. In this experiment described in Full Script 10, a model has been trained to predict ER status on the external dataset first using proliferation and then diagnosis age as an additional input. The results of the parameterized experiment, given in **Table 6**, again showed an increase in the model performance albeit weaker than the previous experiment and proved DeepMed’s ability to generate strong multi-input models.

**Table 6:** Aggregated statistics deployment experiment parameterized to target ER status and histologic neoplasm type given proliferation as an additional input in respective DeepMed runs.

| Additional Input | Target | Class | AUROC | Patient count | p value |
| --- | --- | --- | --- | --- | --- |
| Diagnosis Age | ER Status By IHC | Positive | 0.677 | 68 | 2.41E-02 |
| Diagnosis Age | ER Status By IHC | Negative | 0.677 | 22 | 2.41E-02 |
| Proliferation Score | ER Status By IHC | Positive | 0.872 | 68 | 7.07E-10 |
| Proliferation Score | ER Status By IHC | Negative | 0.872 | 22 | 7.07E-10 |

## Discussion

### Limitations

DeepMed has the capacity to build deep learning networks for patient-level feature prediction directly from histopathology slides. DeepMed is a re-implementation of algorithms that have already been published. ^3,15–17^ The fundamental limitation of this method is that not all clinically significant traits can be predicted from histopathological slides. Two recent large-scale assessments consistently demonstrated that this method predicts approximately one-third of all evaluated genetic changes in human cancer ^4,6^ - and two thirds of all tested molecular alterations are not predictable. However, although this approach is not ubiquitously applicable, it has been shown to provide clinically relevant performance ^15,28^ and can be used to discover previously unknown biological mechanisms ^27,31^. Another restriction is that many non-computer-savvy researchers find obtaining, storing, preparing, and evaluating histology image data difficult. DeepMed intends to reduce the work required to employ deep learning in end-to-end computational pathology by removing this load. However, some of the problems are caused by a lack of standardization in computational pathology (for example, the widespread use of numerous proprietary image file formats) or are inherent to the area (such as the large file size of digitized whole slide images).

### Outlook

DeepMed is a straightforward, scalable, and powerful implementation of end-to-end weakly supervised Deep Learning in histology, as demonstrated here. New technologies, such as multiple instance learning^32^ and vision transformers^12^, have recently been investigated for such weakly supervised prediction challenges. Although there is limited data on these strategies’ real-world performance, some of them may become the de-facto state of the art in the future. DeepMed’s modular design makes it simple to include new technologies without affecting the user experience, performance metrics, or any other high-level aspects of the implementation. As a result, DeepMed could be a versatile and future-proof instrument for academic computational pathology.

## Materials, Methods and Procedure

### Software and hardware requirements and setup

DeepMed has been tested on Windows 10, Windows Server 2019 and Ubuntu 18.04. The first step is to install Python 3.8 on the computer. We recommend using Anaconda (https://www.anaconda.com/). The easiest way to get started is to download the latest version of DeepMed from Github here https://github.com/KatherLab/deepmed and install it with “pip”. On Windows, navigate to the directory containing the code, run the standard terminal or powershell and execute:

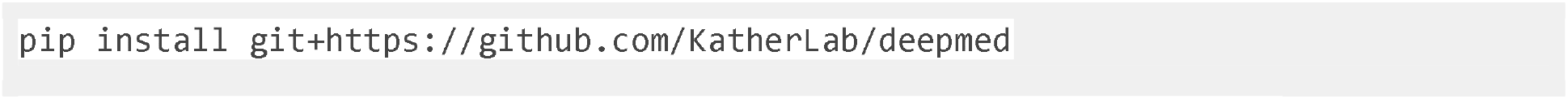

Alternatively, download the codes from Github, navigate to the folder and execute:

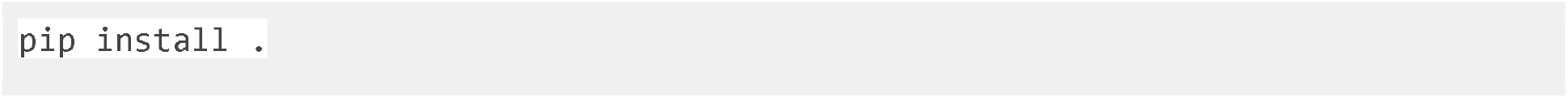

This command will install DeepMed and all its requirements for the current user so that it can be called by other scripts. This will require administrative rights on Windows. Alternatively, the source codes can be directly downloaded from the Github website. DeepMed runs on laptops with an Nvidia graphics processing unit (GPU), on desktop computers with one or multiple Nvidia GPUs or a computing cluster such as EC2 instances on Amazon Web Services (AWS). Throughout this manuscript, we refer to the DeepMed release “v0.8.7” which remains available at https://github.com/KatherLab/deepmed/releases/tag/v0.8.7.

### Data requirements and example data

We have shown the functionalities of DeepMed on two benchmark datasets, TCGA-BRCA-A2 and TCGA-BRCA-E2, that are available at https://zenodo.org/record/5337009^32^. These datasets are already tiled and normalized and were constructed with all images from Walter Reed National Military Medical Center (tissue source site code A2, 100 images) in the TCGA BRCA database and Roswell Park Comprehensive Cancer Center (tissue source site code E2, 90 images) in the TCGA-BRCA database respectively. Here, we use the A2 set for training and the E2 set for testing. For all the experiments herein, the data should be formatted according to the Aachen Protocol ^21^. This requires the image tiles to be situated at the folders named with their parent WSI’s names. Also, for each cohort to be analyzed, there needs to be a clinical table and a slide table. The clinical and slide tables for the benchmark datasets are also available as part of the benchmark dataset. The clinical table has all clinical data that has been saved for each patient including age, sex, cancer type, mutation status etc. Clinical tables include a column which includes patient names and one or multiple columns of target columns (e.g. diagnosis) which includes the label for each patient for the target (e.g. positive or negative). Existing categorical or continuous variables in the clinical table can also be used as inputs in the multi-input models. DeepMed expects the name of the header of the patients column to be “PATIENT” by default in the clinical table. Slide tables, on the other hand, are essential for mapping patients and samples, in this case whole slide image identifiers. They include patient names and corresponding WSI names and, thereby, help DeepMed to ensure that the deep learning method has not been trained and tested on images from the same patient. DeepMed expects the header of the patient and slides columns to be named “PATIENT” and “FILENAME” respectively by default in the slide table.

### Basic workflow

DeepMed allows the user to perform common analyses with very little code. In general, a short Python script (“experiment script”) is enough to run a DeepMed workflow on a dataset which has been prepared in a suitable format. Here, a guideline to construct experiment scripts is given. Apart from the presented functionalities here, all parameters of DeepMed are shown in Suppl. Table 1 and 3.

#### Experiment imports

The first line must be an import statement to import all the necessary functionality. The import statement should be at the top of the experiment script:

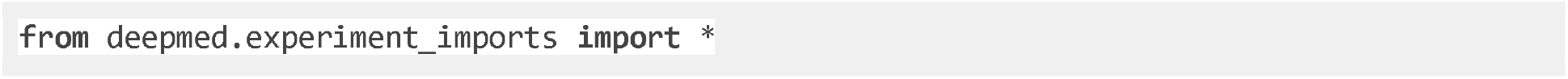

#### Defining the cohorts

In the DeepMed pipeline, both training and deployment is possible to perform on cohorts of patients. Cohorts are defined with the cohort function whose parameters are paths to the directory containing the tiles, the clinical and slide tables, all initalized. ilesDeepMed can also merge multiple cohorts together for both training and testing using `pd.concat()`.

A training cohort consisting of a set of multiple cohorts one to train a neural network model on can be initialized in the following way:

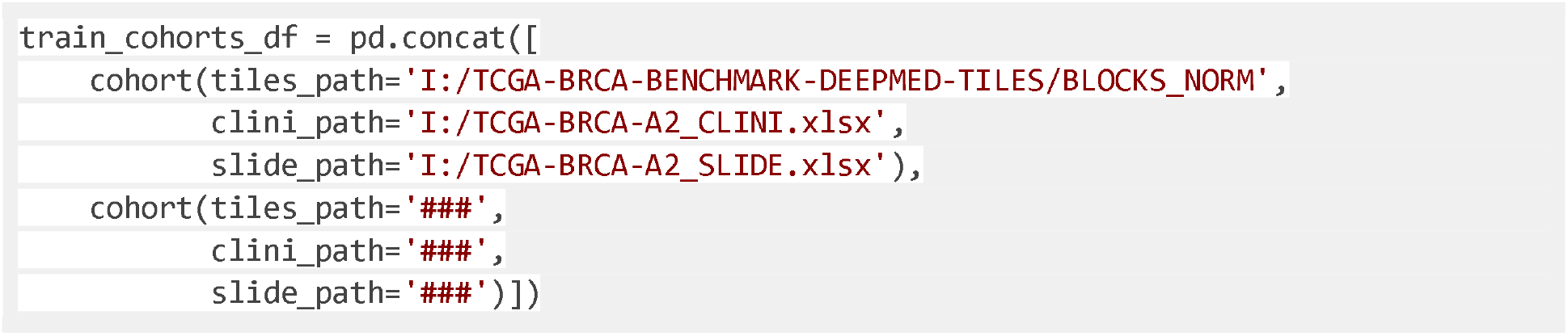

When using Windows-like paths with backslashes, the string for the paths ought to be prefixed with an `r` to prevent the backslashes from being interpreted as character escapes: `tiles_path=r’C:\tile\path’`. The variable `clini_path` holds the string for the path to the clinical data, while `slide_path` indicates the path for the slide table.

#### Defining the experiment structure

Next, the user has to define how to use the selected cohorts. This is done by using TaskGetters, one of the distinguishing features of DeepMed: they define the steps which have to be performed in an experiment. Thanks to their composable structure, they allow the user to easily construct a wide variety of common experiment schemes such as the training and deployment of a singular model, cross-validation, training models on different subgroups or with different hyperparameters. Internally, the TaskGetter will generate a series of tasks, each of which describes a step of our experiment such as preprocessing data, training or deploying a model or evaluating a deployment result. Most of these features are implemented in such a form that they *adapt* other TaskGetters: they take a TaskGetter and modify it. A cross-validation TaskGetter for example would take another TaskGetter and invoke it multiple times, each time with a different training and testing set. Similarly, a subgroup TaskGetter would apply another TaskGetter to one or multiple subsets of our data set. This way, we can nest TaskGetters to build up more and more complex experiment setups.

In the following, we will construct a TaskGetter for a simple, single-target training:

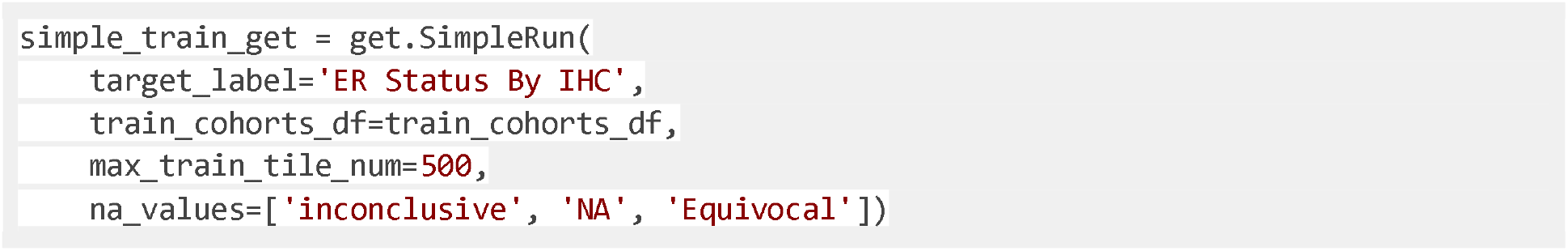

- `get.SimpleRun` describes how to use the data. In this case, we want to train a simple, single-target model. All the following lines describe how this training is to be performed.
- `target_label=‘ER Status By IHC’` dictates the label that the user selects to predict within the model. The clinical table is expected to have a column with that name.
- `train_cohorts_df=train_cohorts_df` are the cohorts to be used for training.
- `max_train_tile_num=500` states how many of a patient’s tiles to sample randomly in the training dataset. Often, increasing the number of tiles for each patient has only a minor effect on the training result. Thus, sampling from a patient’s tiles can significantly speed up training without hugely influencing our results.

- Alternatively, to resample the training tiles used from each slide in each epoch, the option `resample_each_epoch`.
- `na_values=[‘inconclusive’, ‘NA’, ‘Equivocal’]` allows the user to define values which indicate a non-informational training sample. Patients with these indicated labels will be excluded from training.

#### Training the model

For the training to be able to start, the function `do_experiment()` needs to be called in the main script.

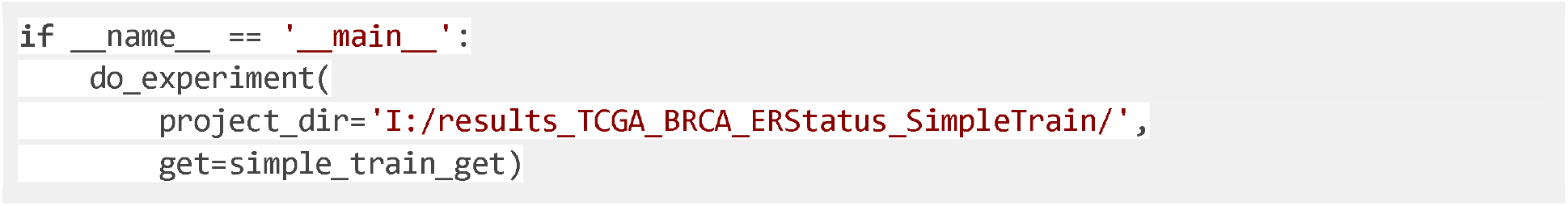

- `project_dir=‘ I:/results_TCGA_BRCA_ERStatus_SimpleTrain/’` defines the location for saving the training results.
- `get=simple_train_get` calls the previously constructed TaskGetter.

#### Deployment (Inference)

Deployment of the neural network model is imperative to ascertain its performance whenever a test dataset is available. In DeepMed, the deployment procedure is quite similar to that of training. After defining the test cohorts, a TaskGetter is constructed with a parameter `test_cohorts_df` instead of `train_cohorts_df:`

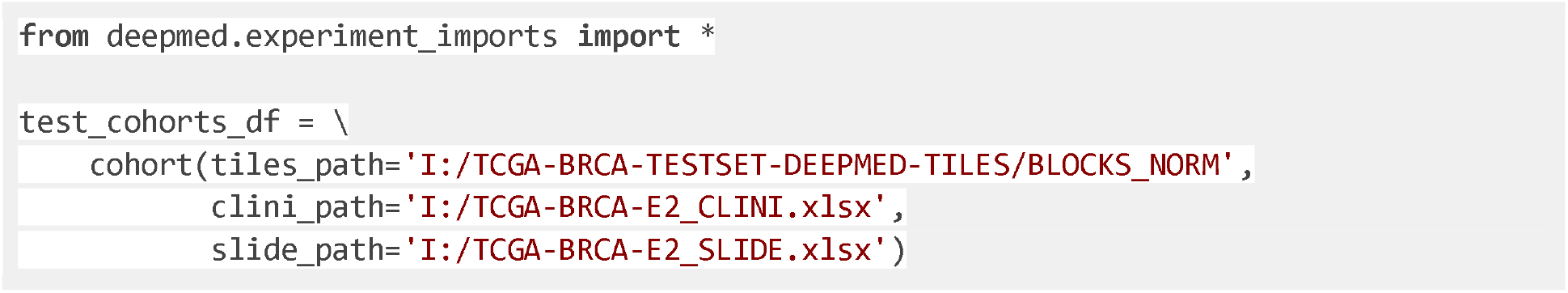

Next, the user has to specify how and from where to load the models to be deployed. Usually, the train parameter inside the TaskGetter is used to further define the modalities of a network’s training. In the case of deployment, a pretrained model is loaded instead of training a model from scratch. The loaded model given to the simple TaskGetter is then deployed at the final step:

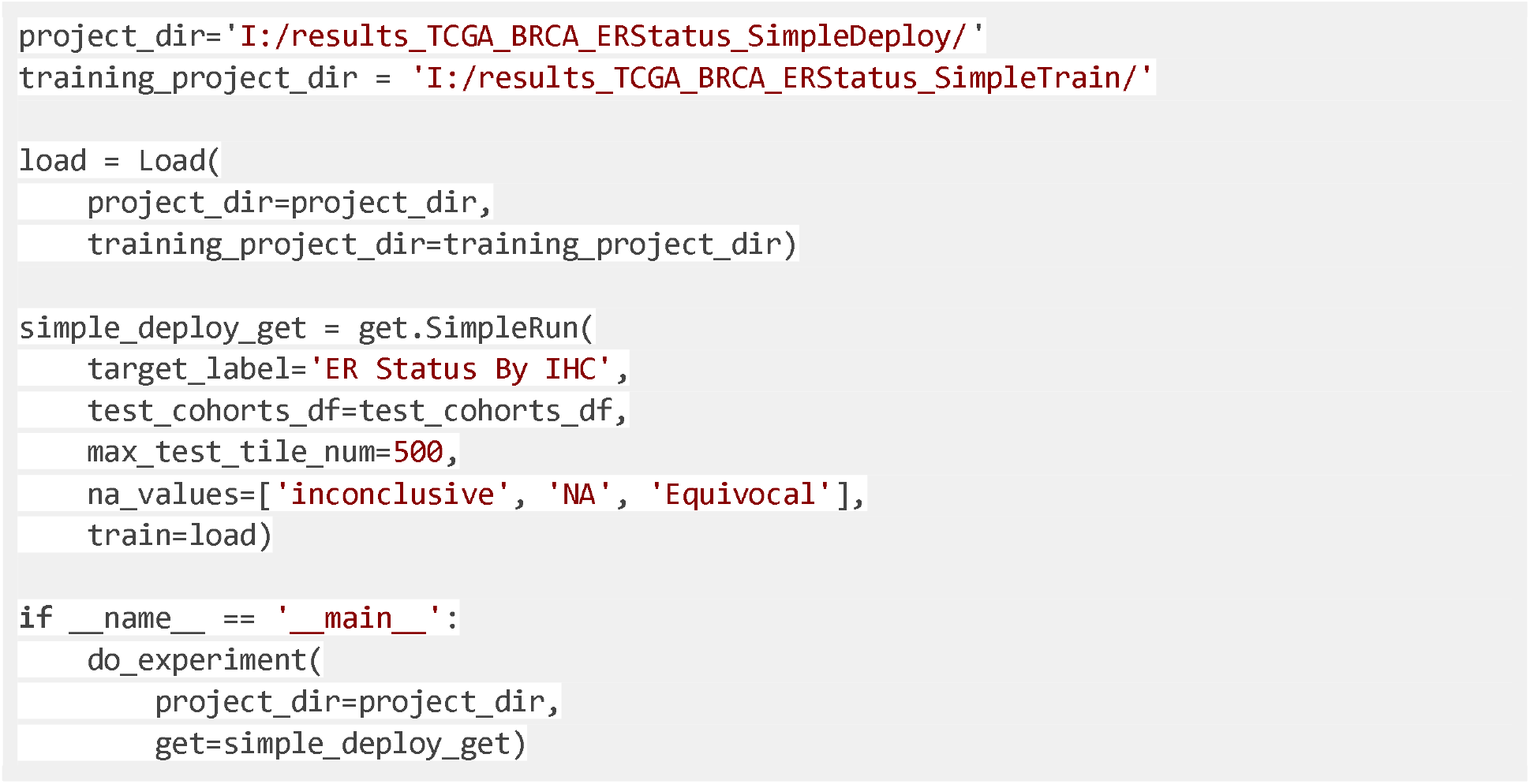

#### Training and Deployment in a single script

With DeepMed, the training and a consecutive deployment are possible to run in the same experiment script. An example for running a training and deployment analysis is given in Full Script 1.

**Full Script 1:**
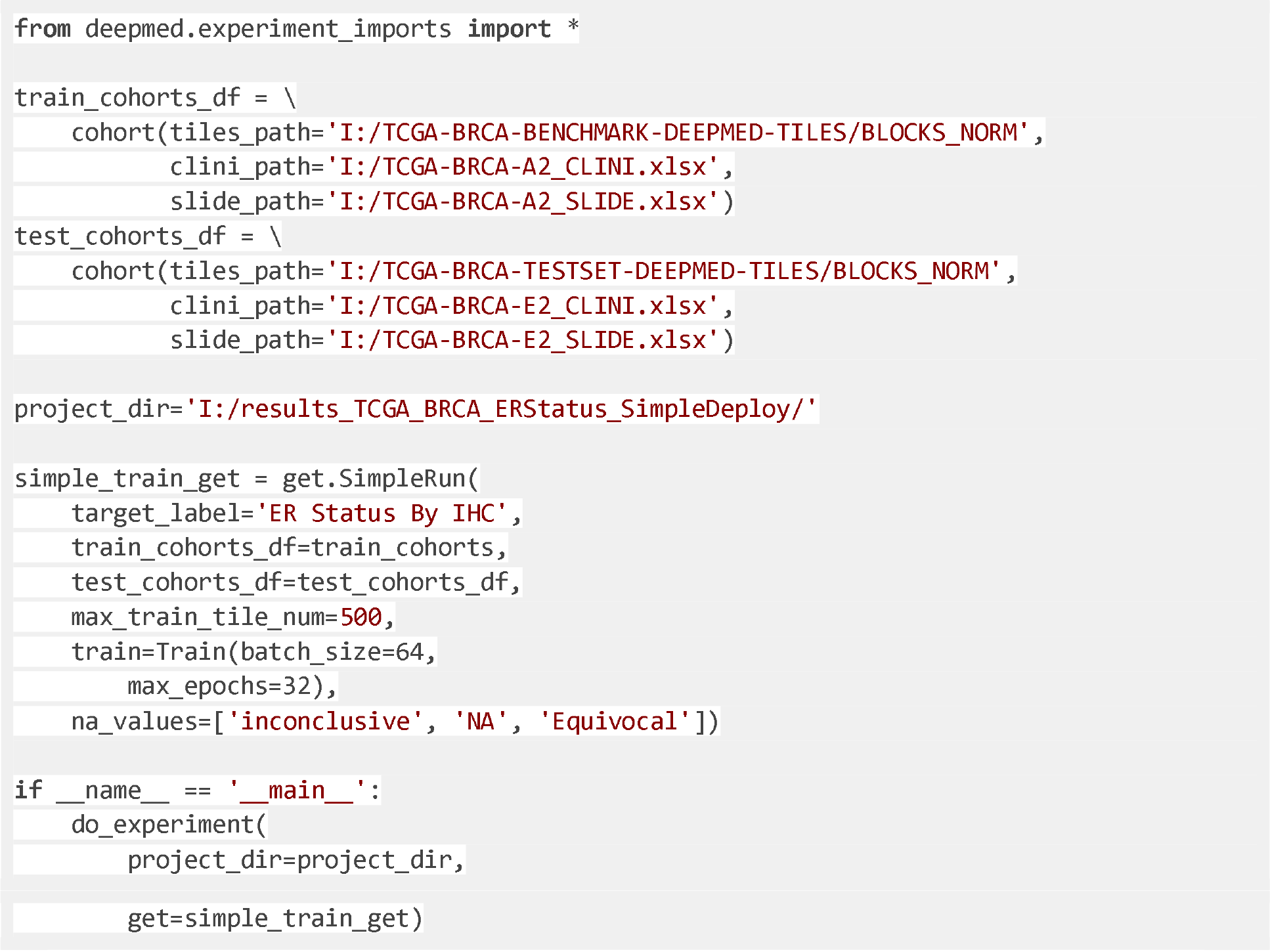
simple_train_and_deploy.py: The experiment script for a simple training.

In Full Script 1, it is shown that the train parameter inside the simple TaskGetter is assigned with the Train function which has the parameters `batch_size` and `max_epochs`. Train is a DeepMed function which describes the details of the single model training. It is an optional function because the used parameters already have the default values, as listed in Supplementary Table 2 along with all other parameters of DeepMed.

#### Defining evaluation Metrics

While the above samples show how to train the model and deploy on a test set, there will not be any statistics or visual output without defining the metrics alongside with the basic parameters. In order to assess the performance of the model on the test set, the parameter evaluators which hold a Python list must be given to the run adapter. For instance:

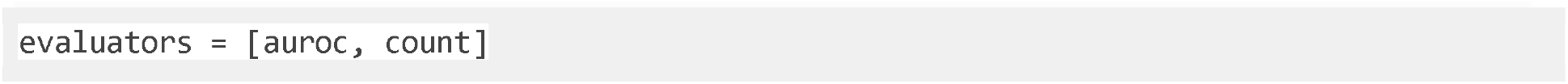

These metrics will calculate the area under the receiver operating characteristic curve (AUROC) and the count of testing samples. However, they are calculated on a tile basis. It is often advantageous to calculate metrics on a per-patient basis instead. This can be done with the Grouped adapter:

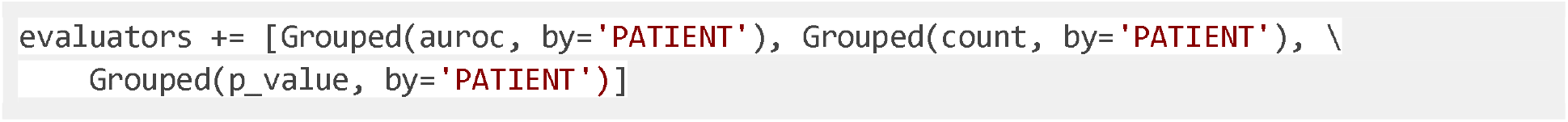

This will modify the AUROC and count metrics in such a way that they are calculated on a per-patient basis instead of a per-tile basis, meaning that instead of the overall tile count per class, the number of patients per class will be calculated. Additionally, p value on a per-patient basis is added into the evaluation metrics, which is calculated by applying a two-tailed t test for differences in the metrics of target classes. Measuring on a per-patient basis is the default behavior of the Group adapter; thus, `by` option can be skipped when grouping is desired on a per-patient basis. The possible evaluation metrics are shown in Suppl. Table 2.

If the deployment script is extended to make use of these evaluators, re-running the script should yield a file called `stats.csv` that contains the requested metrics in the project output directory. The whole experiment script extended with evaluator metrics is given below:

**Full Script 2:**
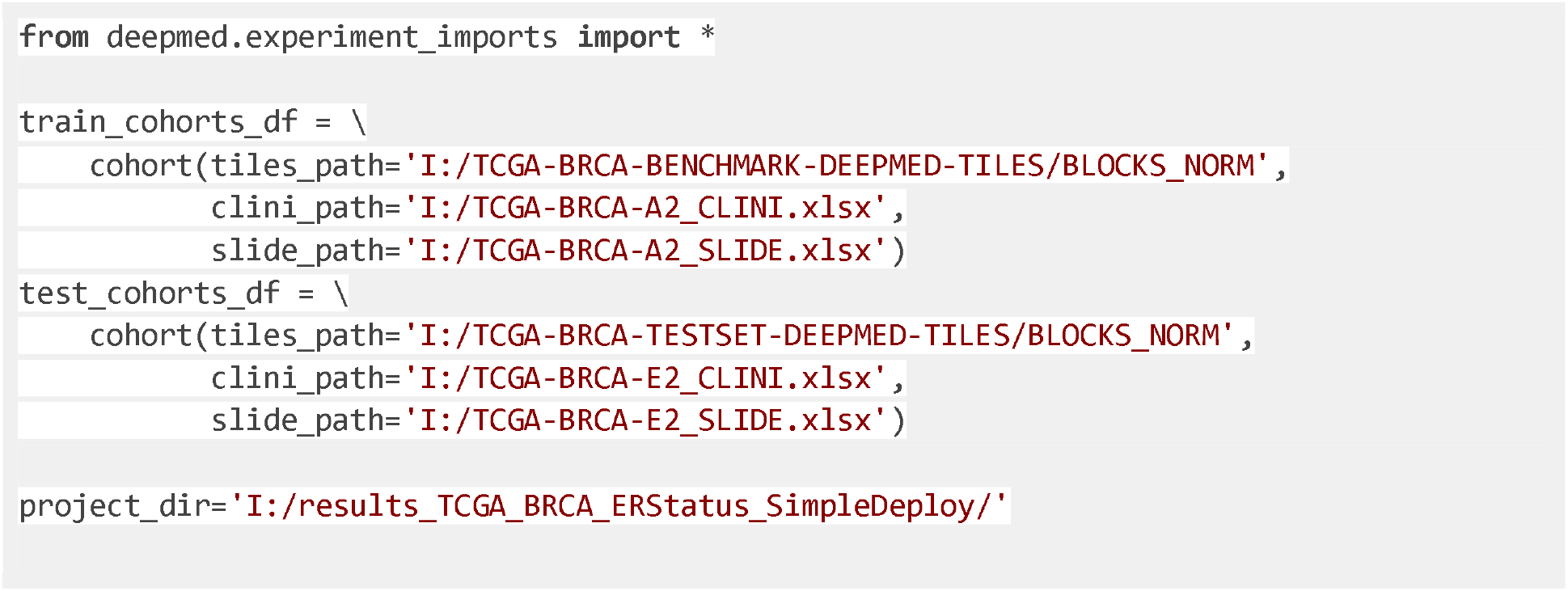
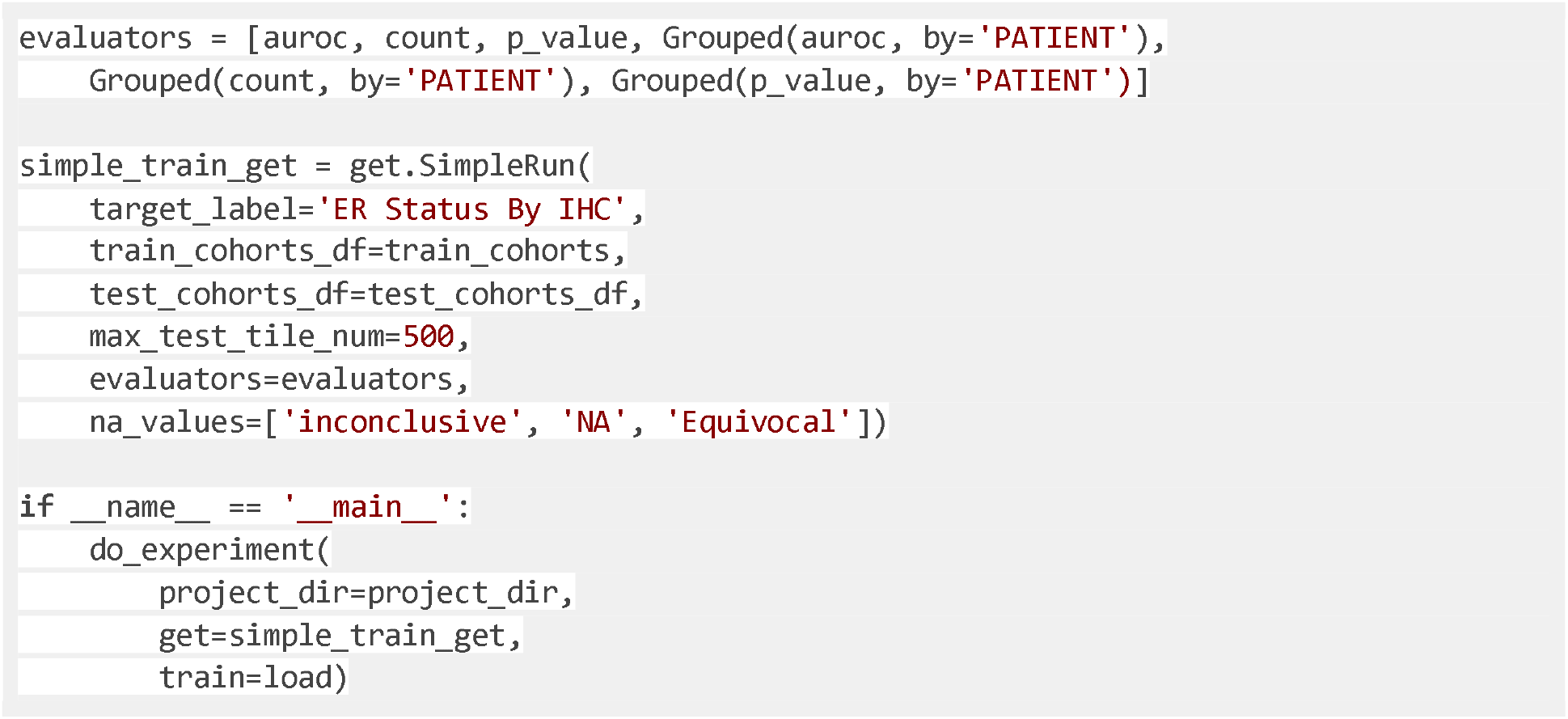
simple_train_and_deploy_w_evaluators.py: The experiment script for a simple training and deployment that has been extended with evaluator metrics.

The presented experiment here sets an example for the *simple run* mode in DeepMed. In other words, the training and deployment tasks with all parameters defined were only on the class-level prediction. Besides *simple run*, DeepMed has several training modes that can be used in any desired combination and order depending on the user’s needs: *multi-target*, *cross-validation*, *subgroup* and *parameterize*. In the following sections, we will introduce the remaining training modes and the data types and parameters that can be used for all training modes.

#### Multi-target training

DeepMed can also run for more than one target with a possibility of using multiple GPUs in the process. A sample script to construct an experiment for a multi-target training is given below with the step by step explanations.

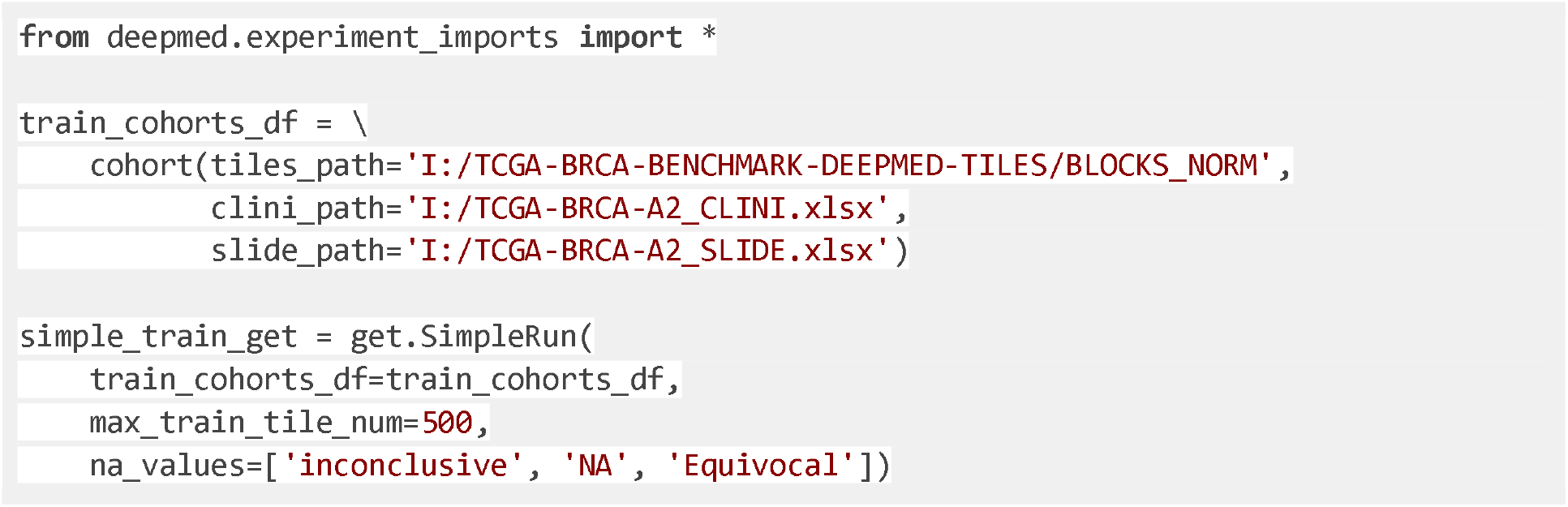

The script starts with a simple TaskGetter as in the previous examples except that the target_label is not specified this time around. The reason for this is to make sure not to restrict the run’s target label, but automatically repeat the training with different target labels. To achieve this, a run adapter is used, which takes another TaskGetter and transforms it instead of generating runs by itself. In this example, a single target TaskGetter will be given and adapted into a multitarget one. How to construct such a run adapter is shown here:

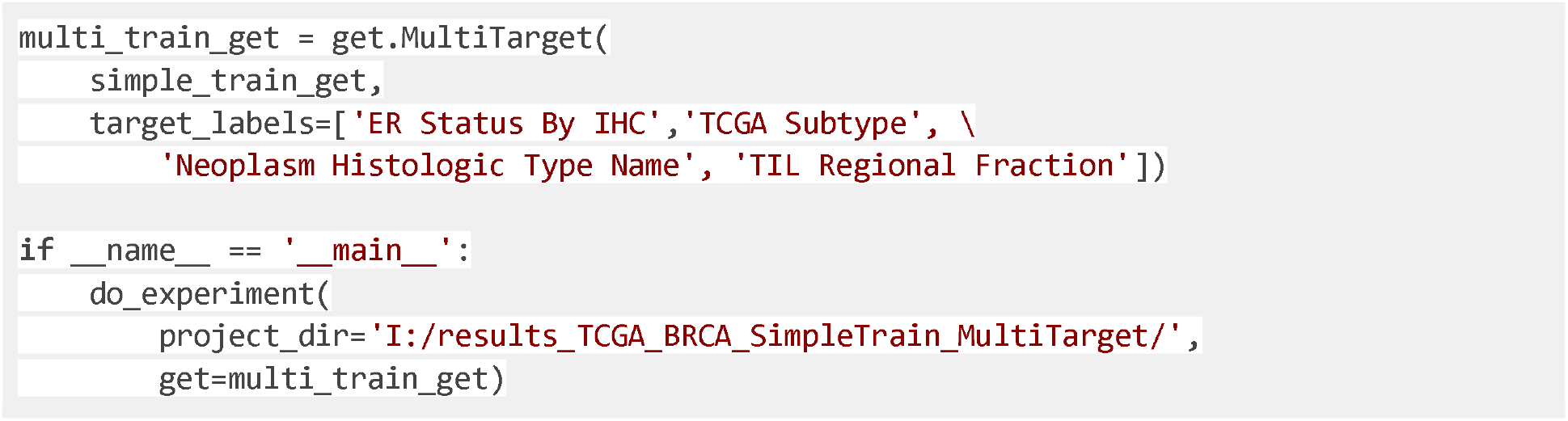

The relative deployment script must be modified for the multiple target data. This is done again with the help of a run adapter that takes the simple deploy TaskGetter as an argument. The deployment script again defines the test cohort and the project directory that is going to store the results. Assuming the model to be deployed is the model that has been trained in the previous example, the training project directory is defined with the output directory of the previously run project that contains the trained model for multiple targets and assigned into the load variable.

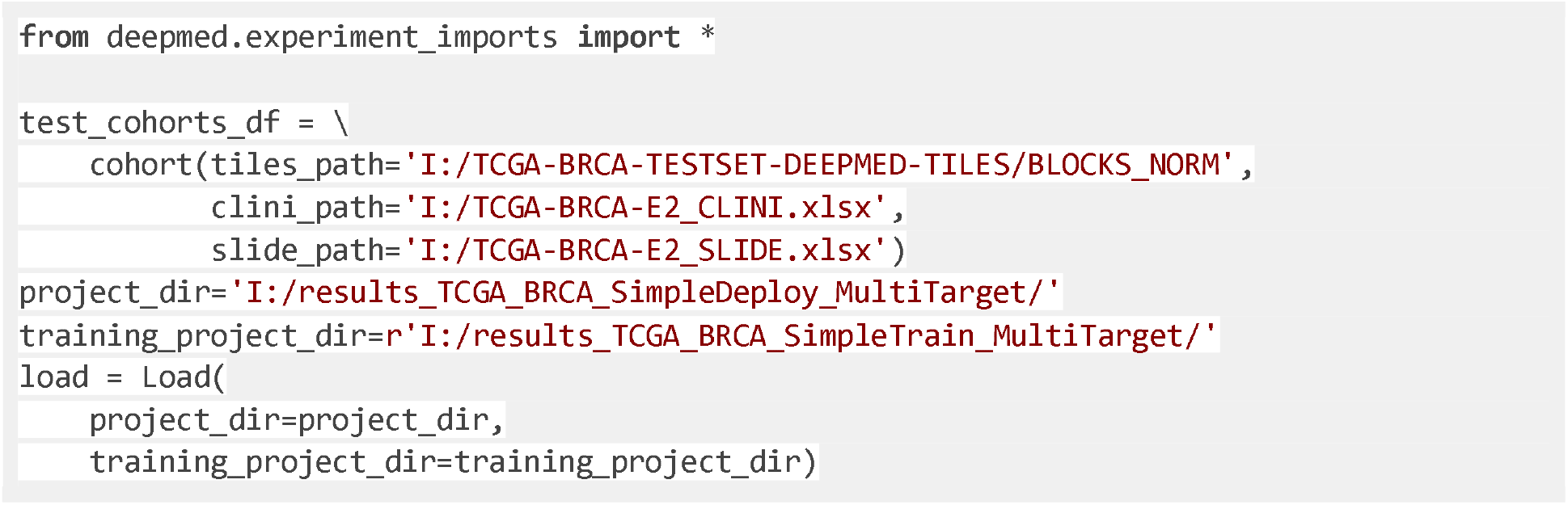

The deploy TaskGetter is able to load the trained model when the `load` variable is passed to the `train` parameter and serves as a template for DeepMed which declares how to perform deployment analysis.

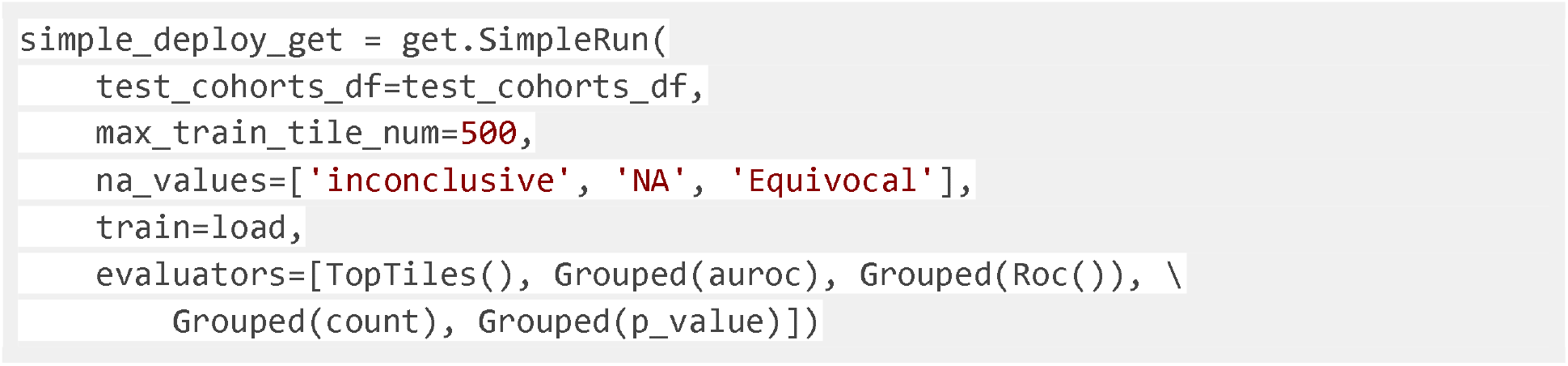

The multi-target TaskGetter takes the simple deploy TaskGetter as an argument, and runs it with its additional functionalities, mainly the multi-target initialization with the parameter `target_labels`. In Full Script 3, multiple targets are set to ER status, TCGA subtype, neoplasm histologic type and TIL regional fraction. The resulting output will be four subdirectories inside the project directory for each target saving the simple run’s predictions, the metrics defined in the simple evaluators and a `stats.csv` file including the results. The `AggregateStats` function in the `multi_target_evaluators` simply concatenates all `stats.csv` files within the subdirectories. The label option is used to give a name to the header of the aggregated column in `stats.csv`.

Therefore, `AggregateStats` can only be used on a higher level TaskGetter than a simple TaskGetter. The `label` parameter writes a user-defined header into the newly created column which is the target names in the concatenated table. The experiment is run when the project directory and the final Task getter is defined in the `do_experiment` function.

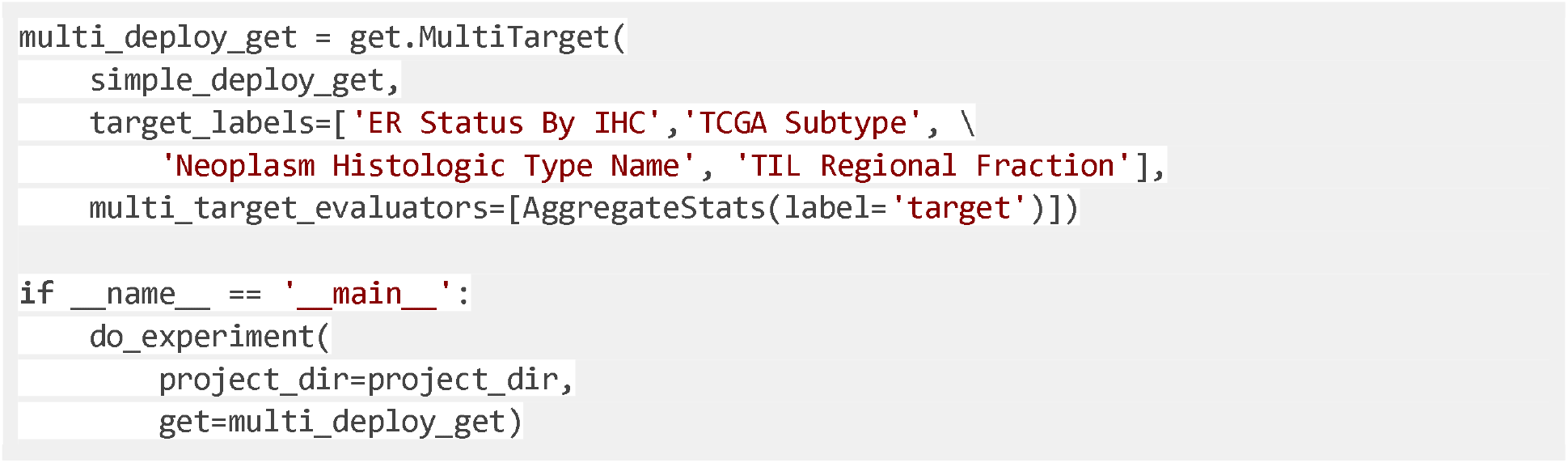

The whole script to deploy a multi-target model is given in Full Script 3.

**Full Script 3:**
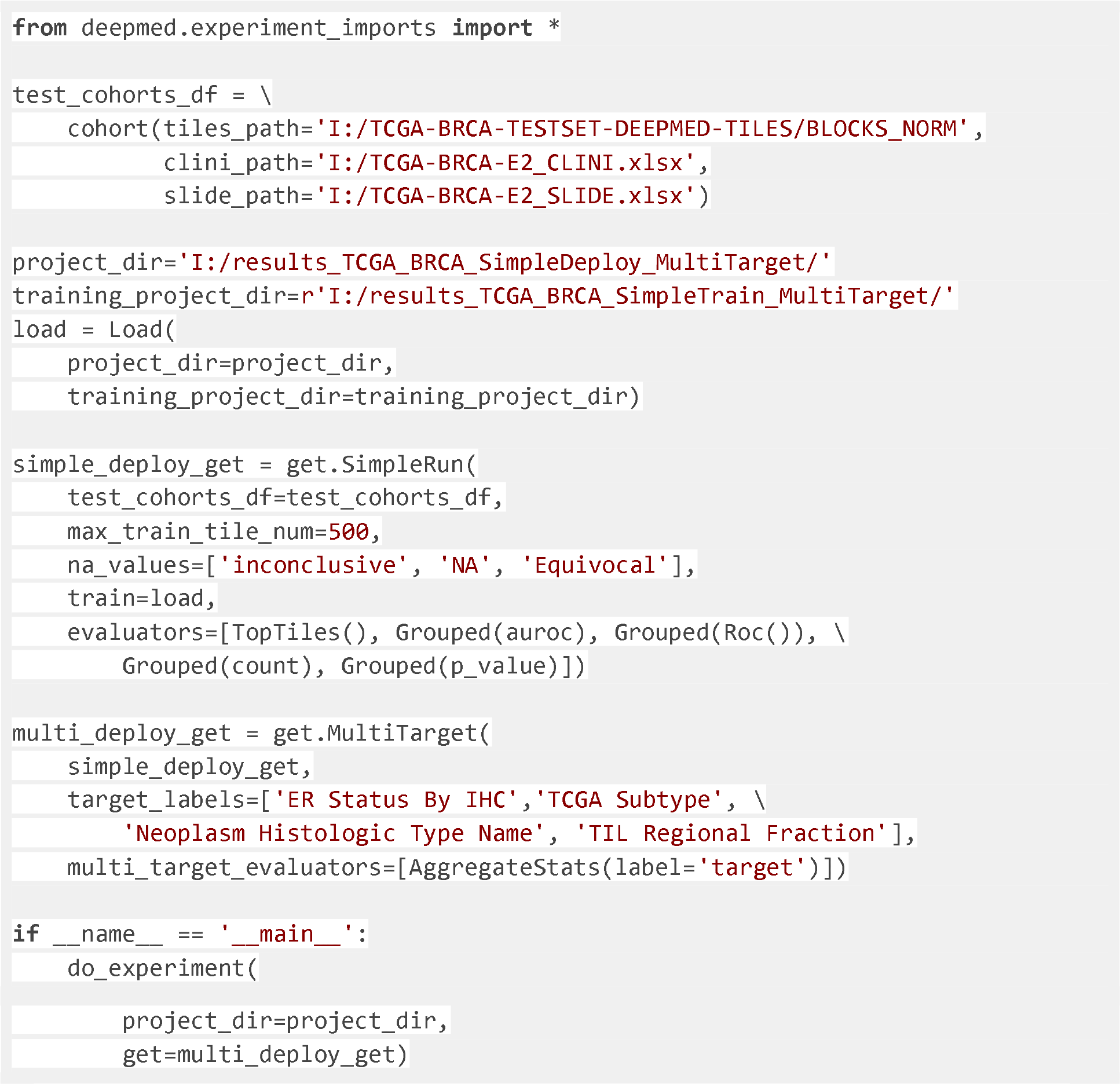
deploy_multitarget.py: The sample experiment script to deploy the trained multi-target model on the benchmark datasets.

It should be noted that it is possible to write a single script to train a model on a training cohort and deploy it on a test cohort in any other configuration of experiment. In this case, to perform the multi-target analysis that has been discussed so far, the user must modify the experiment script in a way that they define training and test cohort and instead of defining a load variable, simply pass the training test cohort to the simple deploy TaskGetter. This is shown in Full Script 4.

**Full Script 4:**
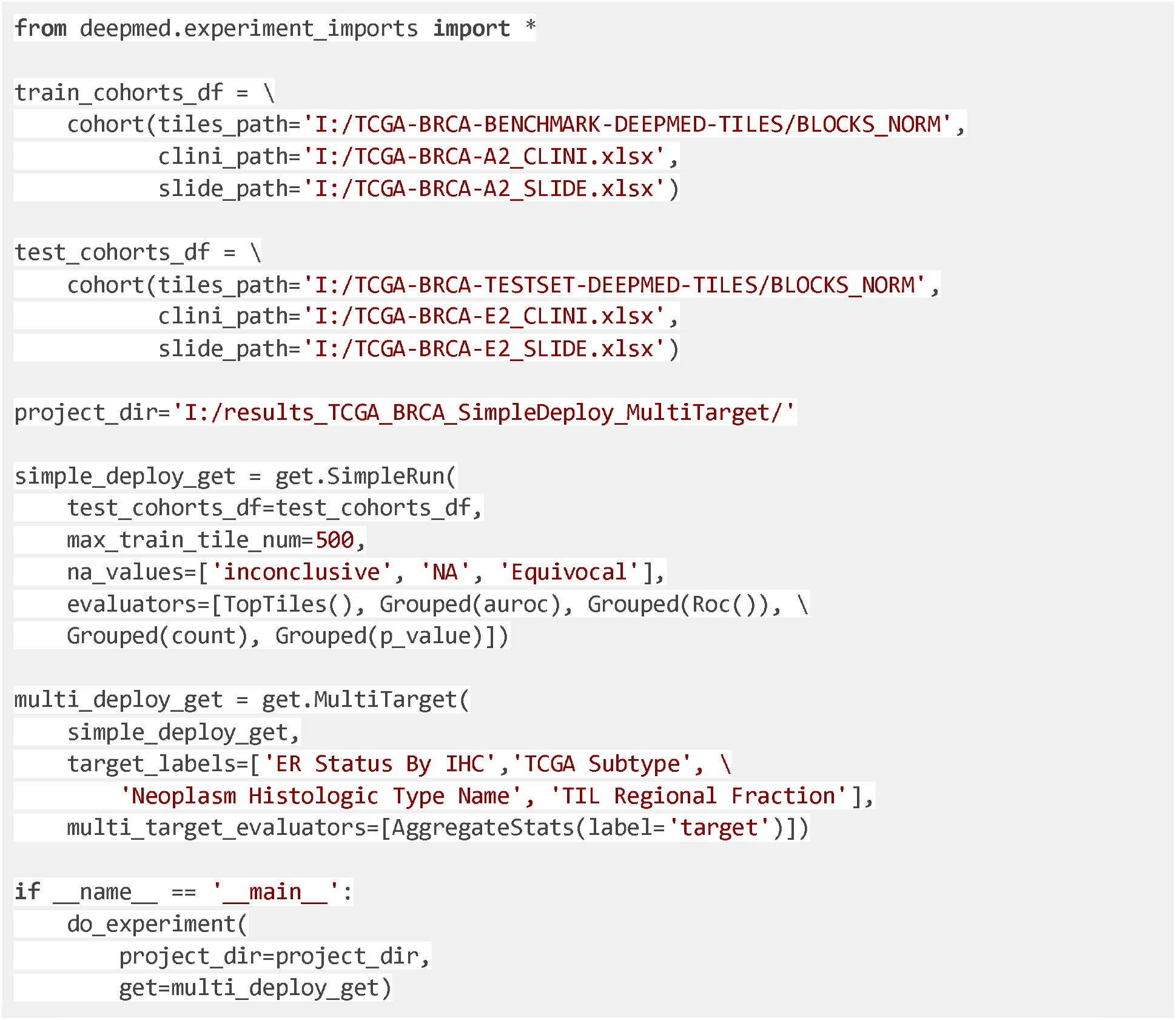
train_and_deploy_multitarget.py: The sample experiment script which includes command to run training and deployment on the benchmark datasets for multiple targets.

### Cross-validation for within-cohort experiments

In addition to simple training, users have the option to perform cross-validation analyses where the training data has been randomly split into training and test datasets for a desired number of times (number of folds), and a model is being trained on each of the different training datasets and evaluated on the test dataset. Cross-validation, in this way, provides users a better overview of performance of the models by having several unseen data by each fold’s models for their evaluation.

A whole sample experiment script to perform a DeepMed analysis with cross-validation is given below in the Full Script 5.

**Full Script 5:**
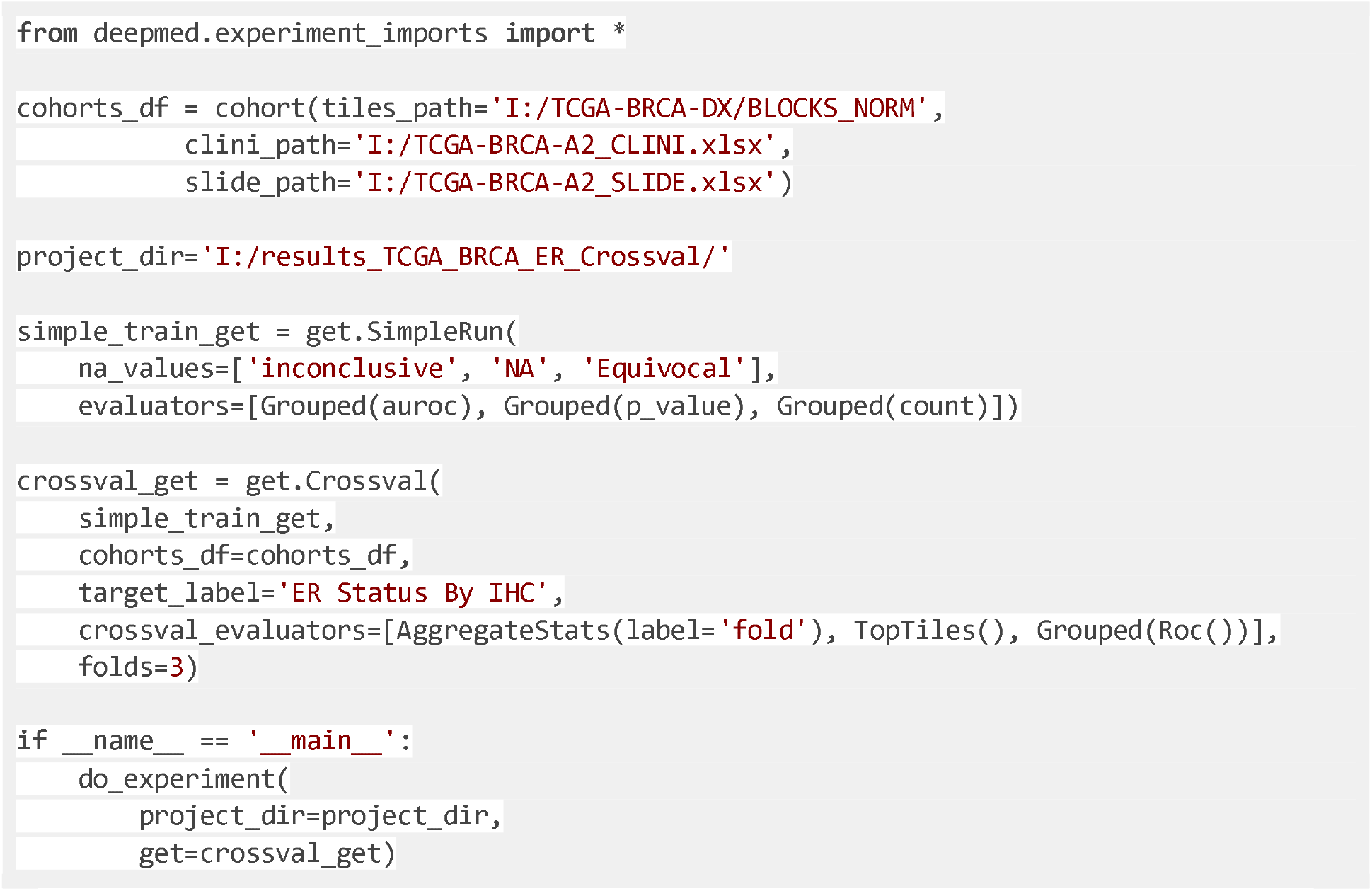
crossvalidated_train.py: The sample experiment script to run within cohort cross-validation analysis to predict ER status only on the benchmark dataset TCGA-BRCA-A2.

In order to run a cross-validated deep learning analysis with DeepMed for a single target, an adapter called Crossval needs to be called. Crossval takes the target label and performs cross validation for it. Once the cross-validation steps have been completed, it applies additional evaluators. These evaluators can then operate on data from all folds. While the evaluators in the single TaskGetter would report AUROC, p value and the patient counts for each model, the evaluators defined in the cross-validation TaskGetter would aggregate these statistics for each target over the different folds and yield top tiles which influenced the model’s decisions at most and the ROC curves on a patient- and fold-level.

When there is more than one target, as seen in the benchmark datasets used here, running a cross validation analysis with DeepMed only requires an additional run adapter for multi-target training. This *multi-target* run adapter executes the cross-validation task getter multiple times, each time with a different target label and output directory, allowing for additional evaluation over all the targets’ cross-validation results. A whole experiment script with the training modes, crossvalidation and multi-target, is given in Full Script 6.

**Full Script 6:**
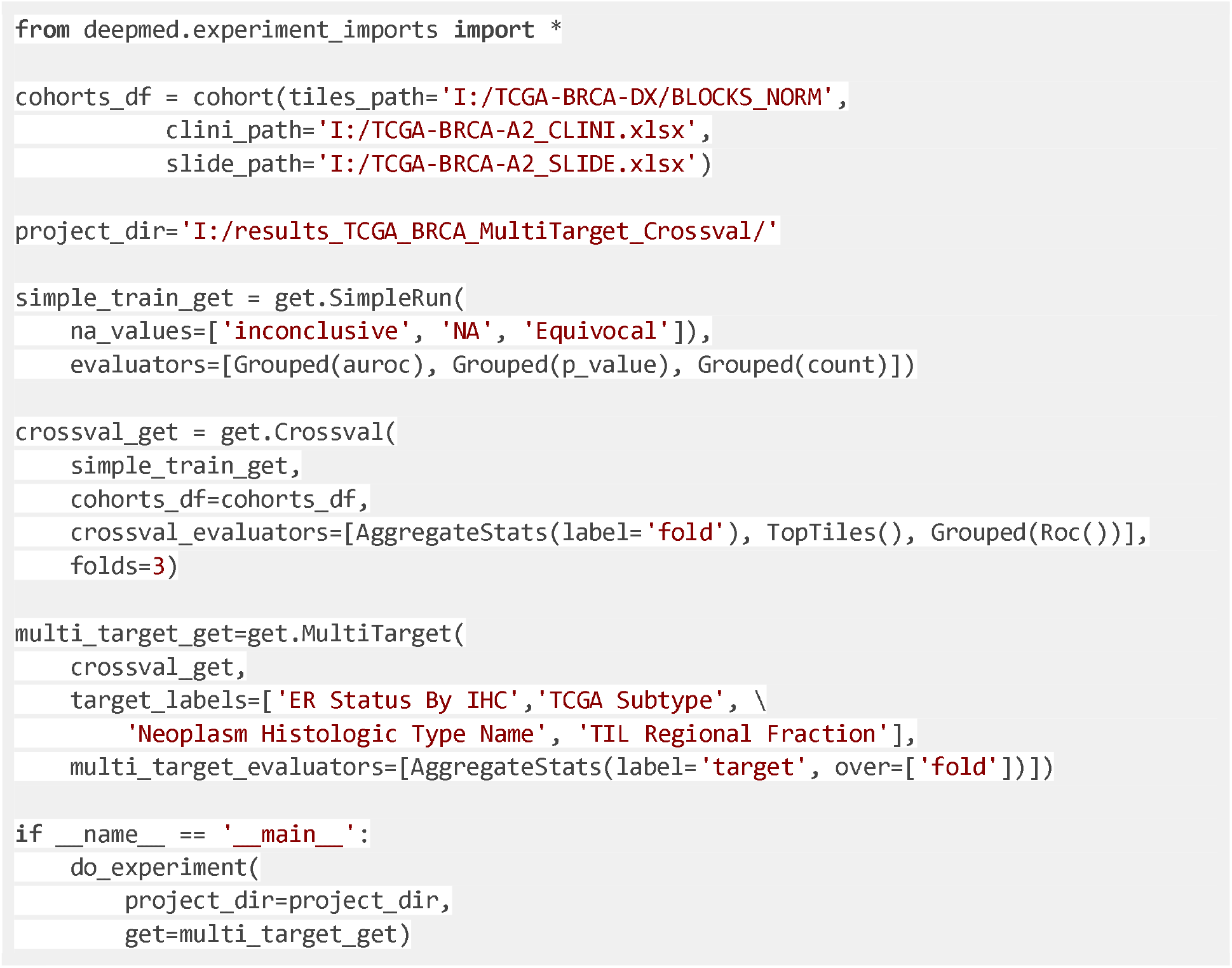
crossvalidated_train_multitarget.py: The sample experiment script to run within-cohort cross-validation analysis on the benchmark dataset TCGA-BRCA-A2.

The Full Script 6 introduces another novelty that is the `over` option in the `AggregateStats`. Without the over option, results from all classes of each target will be reported for all folds in the statistics file. Thus, the over option commands the pipeline to aggregate over the desired column, in this case the folds. It results in a summary of the results including the mean auroc, total patient counts, and p values for the calculations.

### Categorical and continuous targets

In the above samples, the training has been applied to both categorical and continuous data. Categorical targets were “ER Status By IHC” (binary categorical) and “TCGA Subtype” (multiclass categorical). This means the data within the target groups are always a finite number of non-overlapping classes, referring to “Positive” and “Negative” for “ER Status By IHC” and “Basal”, “Lum A” and “Lum B” for “TCGA Subtype”. In addition to categorical data, DeepMed is also able to process and train continuous data where the targets can take any number in an infinite range.

There are two ways that DeepMed handles continuous targets, such as “TMB (nonsynonymous)” (tumor mutation burden): discretization and regression. If the parameter `n_bins` is initialized with a value greater than 0 when the experiment script is handed over to the pipeline, the continuous values are transformed into discrete values by putting them in a desired number of intervals or bins whose borders or cut-off points have been determined by the discretization algorithm. The bins are then subjected to classification in the same way as categorical targets are when being processed. The number of bins is 2 by default, meaning that if the `n_bins` parameter is ignored, the continuous targets are going to be subjected to the binarized classification. If the `n_bins` parameter is set to 0, DeepMed will perform a regression task. To evaluate the performance of such tasks, the coefficient of determination that can be called with `r2` in the evaluators parameter is available in the metrics. Full Script 7 describes an example script to run a regression analysis with a 3-fold cross-validation that is targeting “total number of mutations”.

**Full Script 7:**
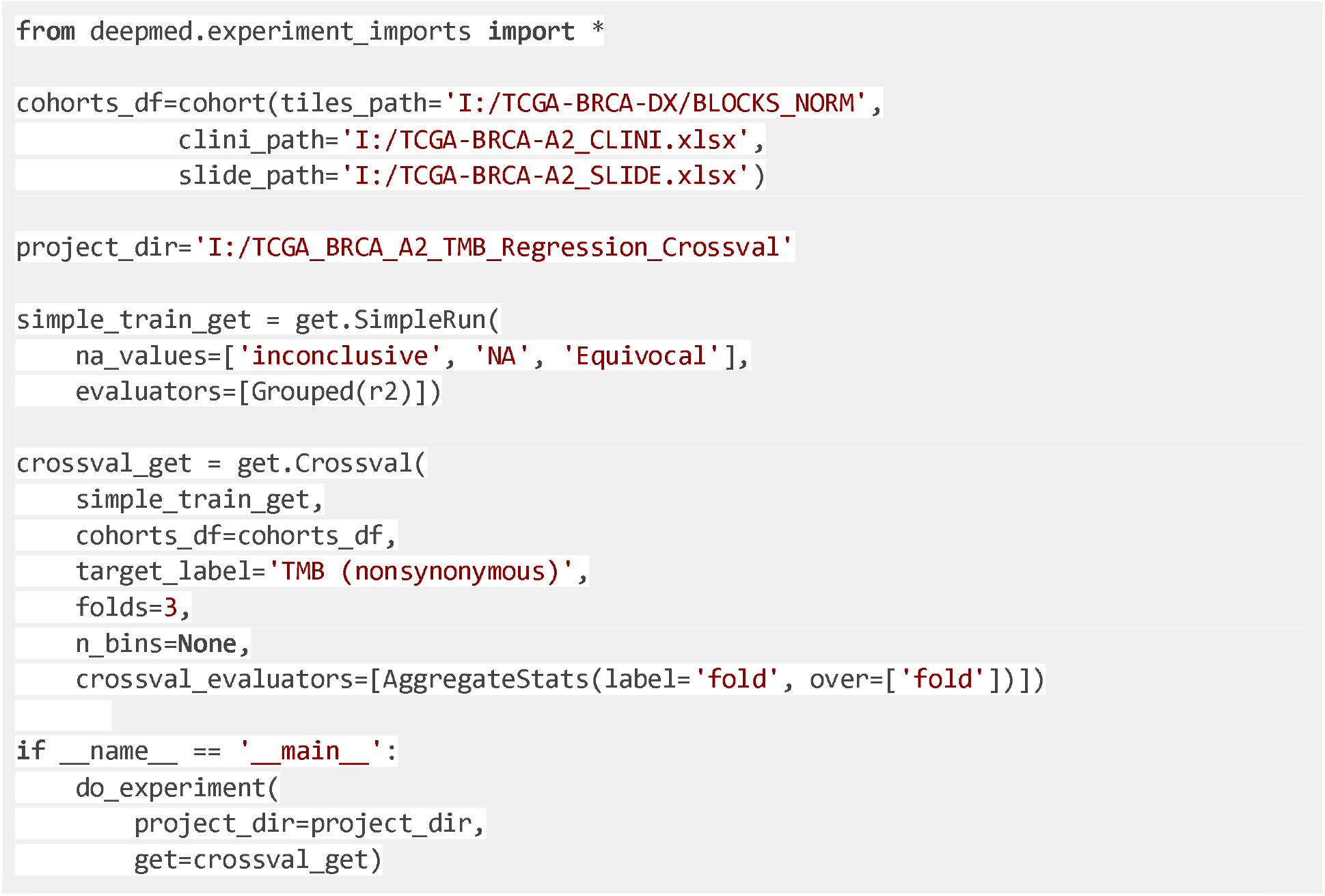
crossvalidated_regression_train_TMB.py: The sample experiment script to run within cohort cross-validation regression analysis on the benchmark dataset TCGA-BRCA-A2.

### Multi-Modality

DeepMed can also perform analysis on data from multiple inputs. These inputs can be of different modalities, for example image or tabular data, clinical or genetic information. In DeepMed, it is possible to provide both continuous and categorical variables to the model as additional inputs. Multiple studies^33,34^ have shown that the performance of Deep Learning models can improve by adding such additional pieces of information to the model. The general practice of combining the modalities is to have a high level embedding of individual modalities is tensor concatenation. In DeepMed, the image input (tiles of whole-slide images) is run through an ImageNet-trained ResNet-18, and the resulting feature vector is concatenated with the chosen tabular data normalized by computing the standard score. The new concatenated network is then given into two fully connected layers and connected to the output layers where predictions are made.

The Full Script 8 shows a whole experiment script to run a multi-modal DeepMed analysis. The `train` parameter within a `SimpleRun` TaskGetter is assigned with the `multi_input.Train` function in order to include the desired tabular information into the training along with image data. The tabular information to be used in the multi-modal training must be present in the clinical table. Both categorical and continuous values can be used as additional inputs by being given to the multi-input training function with the parameters `cats` and `conts` respectively. It is important to note that the names of the additional tabular inputs must always be given inside a Python list to the `cats` and `conts` options.

The Full Script 8 utilizes the multi-modal feature where HER2 status and PR status provide categorical inputs and patient age is provided as a continuous input while training on TCGA-BRCA-A2 dataset and deploying the model on the TCGA-BRCA-E2 dataset in a multi-target mode.

**Full Script 8:**
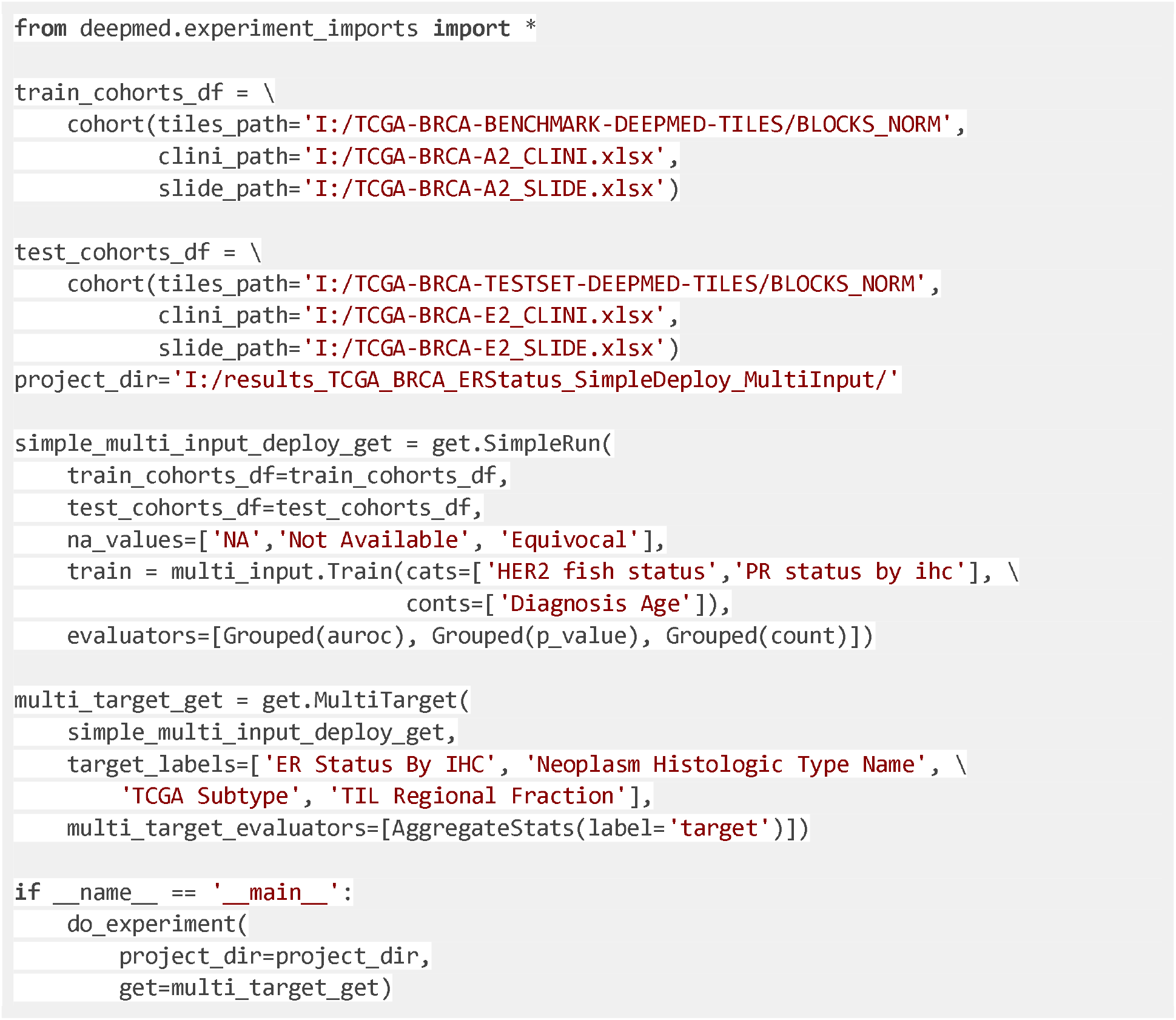
train_and_deploy_multitarget_multiinput.py: The sample experiment script which includes command to run training and deployment with an additional tabular input on the benchmark datasets for multiple targets.

### Subgroup Training

One of the most important training modes of DeepMed is *subgroup training* which allows users to train models for subsets of the original dataset based on user-defined characteristics of the data. This enables us to train different models to analyze subsets of data and compare results. The subgroup training obliges users to create a Python function that describes how to divide the dataset into subsets. Here, we will show how to define a function that would retrieve the subgroups based on TMB values from the clinical table where patients with the TMB value lower than 1.0333 constitute the TMB-low subgroup, and patients a TMB value greater than 1.0333 are analyzed in the TMB-high subgroup. The cut-off value 1.0333, which divides the values under TMB column into 2 bins after dropping NA values, is determined with the *KBinsDiscretizer* function from the preprocessing module of the Python package scikit-learn with a quantile strategy.

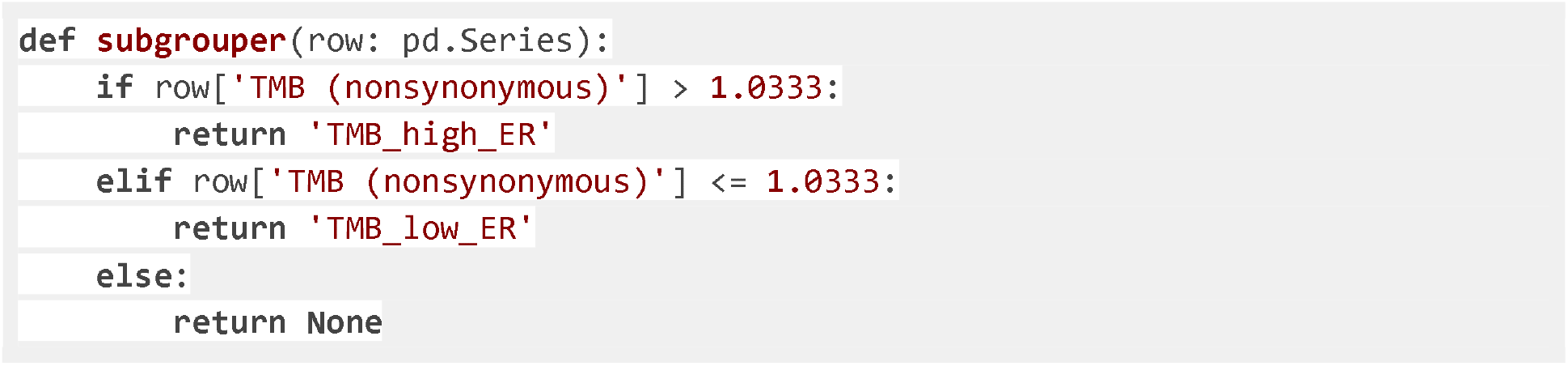

DeepMed takes an input with the data type `pd.Series` because, in the backend of DeepMed, the defined operations described in this function will be applied on each row, which is a pandas (a Python module for data analysis) Series, of the pandas DataFrame.

The Full Script 9 presents a whole DeepMed experiment script to run a simple train-and-deploy experiment to predict the ER status on TMB-low and TMB-high patients separately. Run adapter to pass the simple TaskGetter, in this case, is `Subgroup` TaskGetter. The target label again must be inside the new adapter and the created function’s name must be assigned to the `subgrouper` parameter. The assigned subsets of patients are then used as inputs to train respective models by the subgroup TaskGetter.

**Full Script 9:**
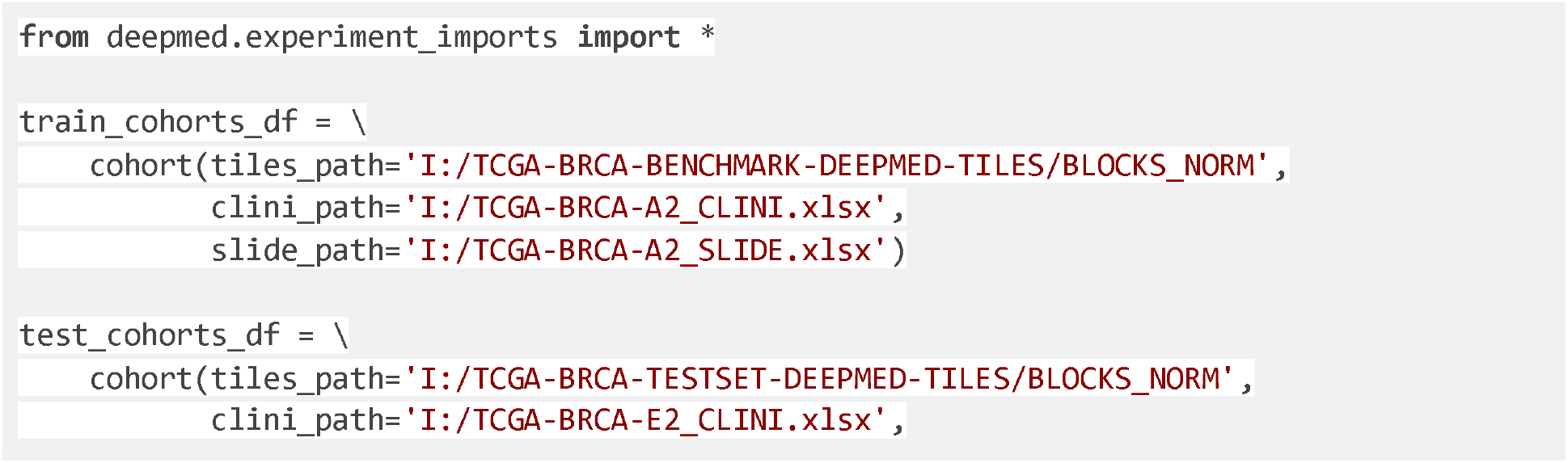
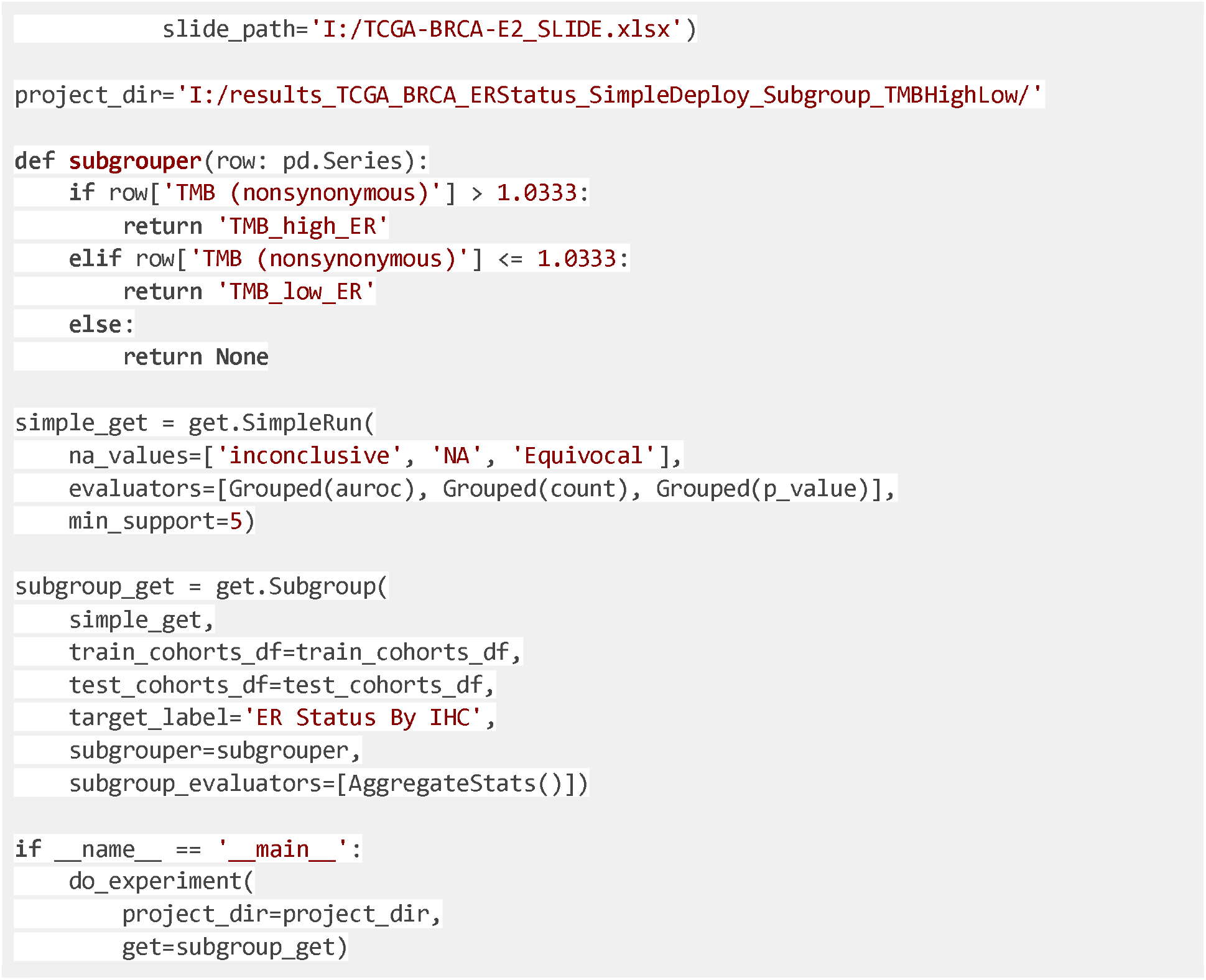
train_and_deploy_subgroup_based_TMB.py: The sample experiment script which runs training and deployment on the subgroups of the benchmark datasets based on their TMB value in order to predict ER status.

The optional parameter `min_support`, that was 10 by default, is set to 2 in Full Script 9. `min_support` parameter dictates the minimum required number of inputs for each target during training. Since there are only 7 samples labeled with negative ER status in the TMB-low subgroup, unless the `min_support` was not lessened, this class is ignored.

### Parameterizing

DeepMed provides users with the opportunity to create and run experiments with different parameters from the same script. This process, called parameterizing, allows the user to run experiments with an unlimited number and combinations of parameters consecutively, thereby saving time and returning aggregated statistics in an orderly fashion.

A parameterization assigns values to arguments which can be given to a TaskGetter. Similarly to the Crossval and MultiTarget adapters, the Parameterize adapter allows us to repeatedly invoke a TaskGetter with different parameterizations. To do so, we supply it with a dictionary which maps the names of the result directories to their respective parameterizations.

In **Full Script 10**, we use parameterization to compare the effects of different auxiliary variables when training multi-modal models. The script will train and evaluate two models, one combining the image data with proliferation status and one combining image data with patient age at diagnosis. The results of these models will be saved in the folders ‘with Proliferation’ and ‘with Diagnosis Age’ respectively. After training both models, it will aggregate these statistics for their results into a single file.

**Full Script 10:**
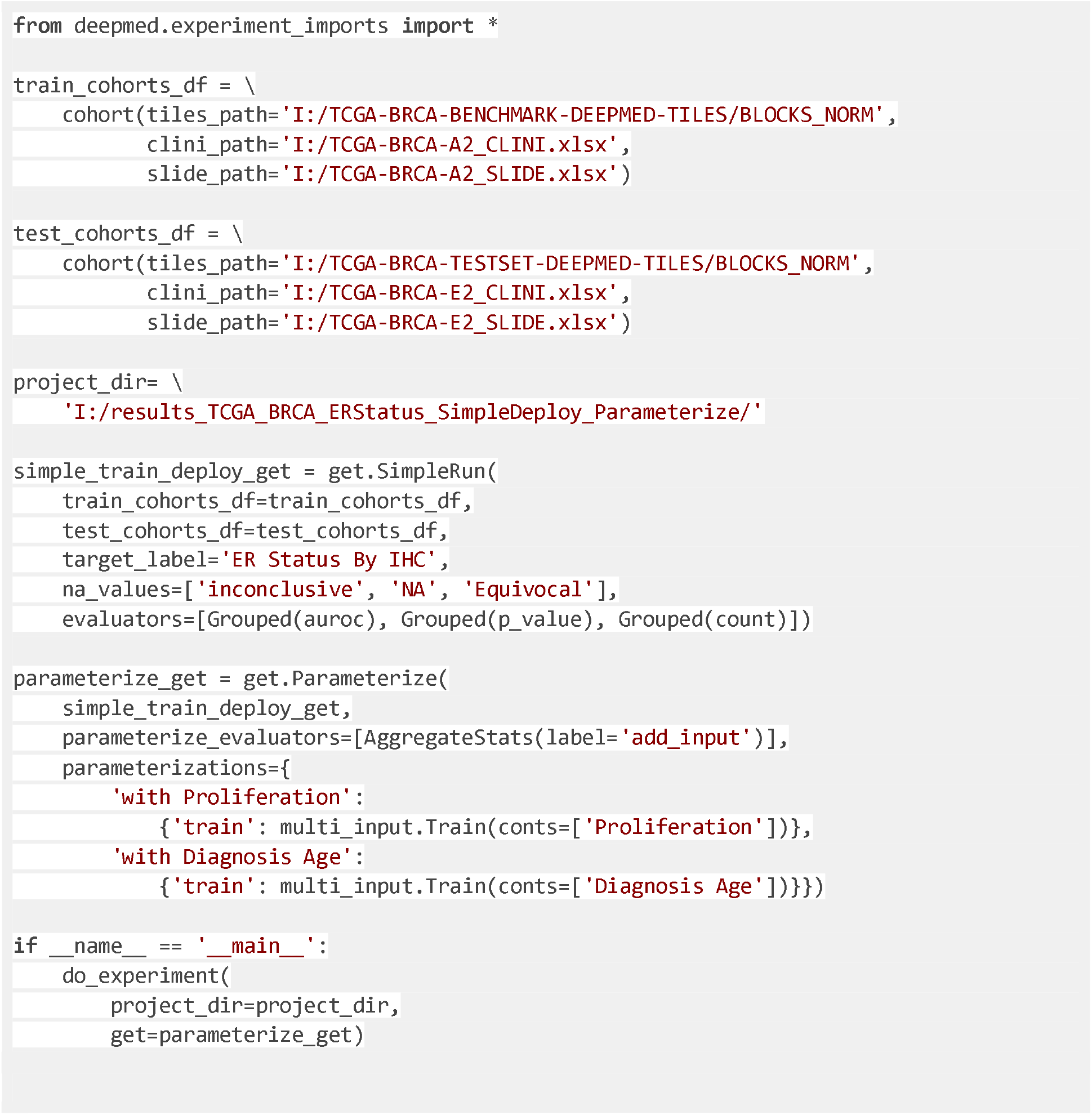
train_and_deploy_parameterizing.py: The sample experiment script which runs training and deployment in a parameterized way using the benchmark datasets.

## Additional information

### Author contributions

MVT, DC, NGL and JNK designed the algorithm. MVT developed the main parts of the software, DC contributed to the software development, conceptualized the manuscript, wrote the manuscript and provided example files. DC, OSL, CL, KJH, HSM, AE and TPS tested the software. All authors contributed to the interpretation of results, writing of the manuscript and collectively made the decision to submit for publication.

## References

1. Calderaro, J. & Kather, J. N. Artificial intelligence-based pathology for gastrointestinal and hepatobiliary cancers. Gut gutjnl–2020 (2020) doi:10.1136/gutjnl-2020-322880.

2. Echle, A. et al. Deep learning in cancer pathology: a new generation of clinical biomarkers. British Journal of Cancer (2020) doi:10.1038/s41416-020-01122-x.

3. Kather, J. N. et al. Deep learning can predict microsatellite instability directly from histology in gastrointestinal cancer. Nat. Med. 25, 1054–1056 (2019).

4. Kather, J. N. et al. Pan-cancer image-based detection of clinically actionable genetic alterations. Nature Cancer 1, 789–799 (2020).

5. Schmauch, B. et al. A deep learning model to predict RNA-Seq expression of tumours from whole slide images. Nature Communications vol. 11 (2020).

6. Fu, Y. et al. Pan-cancer computational histopathology reveals mutations, tumor composition and prognosis. Nature Cancer 1–11 (2020).

7. Saillard, C. et al. Predicting survival after hepatocellular carcinoma resection using deeplearning on histological slides. Hepatology (2020) doi:10.1002/hep.31207.

8. Bychkov, D. et al. Deep learning identifies morphological features in breast cancer predictive of cancer ERBB2 status and trastuzumab treatment efficacy. Sci. Rep. 11, 4037 (2021).

9. Coudray, N. et al. Classification and mutation prediction from non–small cell lung cancer histopathology images using deep learning. Nat. Med. 24, 1559–1567 (2018).

10. Woerl, A.-C. et al. Deep Learning Predicts Molecular Subtype of Muscle-invasive Bladder Cancer from Conventional Histopathological Slides. Eur. Urol. 78, 256–264 (2020).

11. Binder, A. et al. Morphological and molecular breast cancer profiling through explainable machine learning. Nature Machine Intelligence 1–12 (2021).

12. Laleh, N. G. et al. Benchmarking artificial intelligence methods for end-to-end computational pathology. bioRxiv 2021.08.09.455633 (2021) doi:10.1101/2021.08.09.455633.

13. Foersch, S. et al. Deep learning for diagnosis and survival prediction in soft tissue sarcoma. Ann. Oncol. (2021) doi:10.1016/j.annonc.2021.06.007.

14. Lu, M. Y. et al. Data-efficient and weakly supervised computational pathology on whole-slide images. Nature Biomedical Engineering 1–16 (2021).

15. Echle, A. et al. Clinical-Grade Detection of Microsatellite Instability in Colorectal Tumors by Deep Learning. Gastroenterology 159, 1406–1416.e11 (2020).

16. Muti, H. S. et al. Development and validation of deep learning classifiers to detect Epstein-Barr virus and microsatellite instability status in gastric cancer: a retrospective multicentre cohort study. The Lancet Digital Health 0, (2021).

17. Loeffler, C. M. L. et al. Artificial Intelligence–based Detection of FGFR3 Mutational Status Directly from Routine Histology in Bladder Cancer: A Possible Preselection for Molecular Testing? European Urology Focus (2021) doi:10.1016/j.euf.2021.04.007.

18. Schrammen, P. L. et al. Weakly supervised annotation free cancer detection and prediction of genotype in routine histopathology. The Journal of Pathology (2021) doi:10.1002/path.5800.

19. Ghaffari Laleh, N. et al. Deep Learning for interpretable end-to-end survival (E-ESurv) prediction in gastrointestinal cancer histopathology. in Proceedings of the MICCAI Workshop on Computational Pathology (eds. Atzori, M. et al.) vol. 156 81–93 (PMLR, 2021).

20. Kiehl, L. et al. Deep learning can predict lymph node status directly from histology in colorectal cancer. Eur. J. Cancer 157, 464–473 (2021).

21. Muti, H. S. et al. The Aachen Protocol for Deep Learning Histopathology: A hands-on guide for data preprocessing. (2020) doi:10.5281/ZENODO.3694994.

22. Macenko, M. et al. A method for normalizing histology slides for quantitative analysis. in 2009 IEEE International Symposium on Biomedical Imaging: From Nano to Macro 1107–1110 (2009).

23. Bankhead, P. et al. QuPath: Open source software for digital pathology image analysis. Sci. Rep. 7, 16878 (2017).

24. Krizhevsky, A., Sutskever, I. & Hinton, G. E. ImageNet Classification with Deep Convolutional Neural Networks. in Advances in Neural Information Processing Systems 25 (eds. Pereira, F., Burges, C. J. C., Bottou, L. & Weinberger, K. Q.) 1097–1105 (Curran Associates, Inc., 2012).

25. Couture, H. D. et al. Image analysis with deep learning to predict breast cancer grade, ER status, histologic subtype, and intrinsic subtype. NPJ Breast Cancer 4, 30 (2018).

26. Naik, N. et al. Deep learning-enabled breast cancer hormonal receptor status determination from base-level H&E stains. Nat. Commun. 11, 5727 (2020).

27. Brockmoeller, S. et al. Deep Learning identifies inflamed fat as a risk factor for lymph node metastasis in early colorectal cancer. J. Pathol. (2021) doi:10.1002/path.5831.

28. Campanella, G. et al. Clinical-grade computational pathology using weakly supervised deep learning on whole slide images. Nat. Med. 25, 1301–1309 (2019).

29. Skrede, O.-J. et al. Deep learning for prediction of colorectal cancer outcome: a discovery and validation study. Lancet 395, 350–360 (2020).

30. Schulz, S. et al. Multimodal deep learning for prognosis prediction in renal cancer. Front. Oncol. 11, 788740 (2021).

31. Wulczyn, E. et al. Interpretable Survival Prediction for Colorectal Cancer using Deep Learning. arXiv [eess.IV] (2020).

32. Kather, J. N. Histopathology images for end-to-end AI, based on TCGA-BRCA. (2021). doi:10.5281/zenodo.5337009.

33. Höhn, J. et al. Combining CNN-based histologic whole slide image analysis and patient data to improve skin cancer classification. Eur. J. Cancer 149, 94–101 (2021).

34. Chen, R. J. et al. Pathomic Fusion: An Integrated Framework for Fusing Histopathology and Genomic Features for Cancer Diagnosis and Prognosis. IEEE Transactions on Medical Imaging 1–1 (2020) doi:10.1109/tmi.2020.3021387.

