## Supplementary material for "DeepMed: A unified, modular pipeline for end-to-end deep learning in computational pathology": Suppl. Table 1-2

### Supplementary Tables

| **Parameter** | **Data type** | **Default value** | **Description** |
| --- | --- | --- | --- |
| **Main** | | | |
| **Defining experiment (**do_experiment()**)** | | | |
| project_dir | String or path | - | Directory to save project data in. |
| get | TaskGetter | - | A function which generates tasks. |
| num_concurrent_tasks | String | None | Maximum amount of tasks to be run at the same time. If None, the number of tasks will grow with the number of available devices. If 0, all jobs will be run in the main process (useful for debugging). |
| devices | Map type* | {0: 4} | Devices to use for training and the maximum number or models to be trained at once for each device |
| logfile | String | “logfile” | Name of the log file. |
| keep_going | Boolean | False | Whether to stop all runs on an exception. |
| **Defining Cohort (**cohort()**)** | | | |
| tiles_path | String or path | - | Path where image tiles are stored. All tiles have to be stored in a directory named after the original slide. These directories are located under the directory specified by the "tiles_path" |
| clini_path | String or path | - | Path to the clinical table, which maps a patient identifier ("patient_label") to clinical information, either in csv or excel format. |
| slide_path | String or path | - | Path to the slide table, which maps the slide identifier ("slidename_label") to the patient identifier ("patient_label"), either in csv or excel format. |
| patient_label | String | “PATIENT” | Column of clinical and slide tables containing patient IDs. |
| slidename_label | String | “FILENAME” | Column of the slide table containing the slide names. |
| **TaskGetters** | | | |
| **SimpleRun()** | | | |
| target_label | String | - | Column name of the target to be predicted from the clinical table. |
| train_cohorts_df | Cohort (pd.DataFrame) | None | Cohort to use for training. |
| test_cohorts_df | Cohort (pd.DataFrame) | None | Cohort to test on. |
| patient_label | String | “PATIENT” | Column of clinical and slide tables containing patient IDs. |
| balance | Boolean | True | Whether the training set should be balanced. Applies to categorical targets only. |
| train | Callable | Train() | A function training a model, e.g. the function objects returned by Train(), multi_input.Train() etc. |
| resample_each_epoch | Boolean | False | Whether to resample the training tiles used from each slide each epoch. |
| max_train_tile_num | Integer | 128 | The maximum number of tiles per patient to use for training in each epoch. |
| max_valid_tile_num | Integer | 256 | Maximum number of validation tiles used in each epoch. |
| max_test_tile_num | Integer | 512 | Maximum number of testing tiles used in each epoch. |
| valid_frac | Float | 0.2 | Fraction of patients which will be reserved for validation during training. |
| n_bins | Integer | 2 | Number of bins to discretize continuous values into. |
| na_values | Iterable of any data type | [ ] | Class labels whose sample to ignore during training. |
| min_support | Integer | 10 | The minimum amount of class samples required for the class to be included in training. Classes with less support are dropped. |
| evaluators | Iterable of Evaluator class | [ ] | List of evaluation metrics. |
| max_class_count | Map type* | None | A dictionary mapping class name to desired number of patients to be analyzed. |
| **MultiTarget()** | | | |
| get | TaskGetter | - | TaskGetter to adapt. |
| target_labels | Iterable of string type | - | Target labels to invoke “get” on. |
| multi_target_evaluators | Iterable of evaluators | [ ] | List of evaluation metrics. |
| **Crossval()** | | | |
| get | TaskGetter | - | TaskGetter to adapt. |
| target_label | String | - | Target column name to be predicted from the clinical table. |
| cohorts_df | Cohort (pd.DataFrame) | - | Cohort to perform cross-validated analysis on. |
| folds | Integer | 3 | Number of folds in cross-validation. |
| n_bins | Integer | 2 | Number of bins to discretize continuous values into. |
| na_values | Iterable of any data type | [ ] | Class labels whose sample to ignore during training. |
| min_support | Integer | 10 | Minimum amount of class samples required for the class per fold to be included in training. Classes with less support are dropped. |
| patient_label | String | “PATIENT” | Column of clinical and slide tables containing patient IDs. |
| crossval_evaluators | Iterable of evaluators | [ ] | List of evaluation metrics. |
| **Subgroup()** | | | |
| get | TaskGetter | - | TaskGetter to adapt. |
| target_label | String | - | Target column name to be predicted from the clinical table. |
| subgrouper | Callable | - | A function mapping a sample of the training dataset into a subgroup. The function is given a row from the training dataset and has to return either a string describing the group name, or None if it is to be excluded from training. |
| subgroup_evaluators | Iterable of evaluators | [ ] | A list of evaluators to be executed after all subgroup runs have been completed. |
| **Parameterize()** | | | |
| get | TaskGetter | - | TaskGetter to adapt. |
| parameterizations | Map type* | - | A dictionary mapping names to parameterizations. |
| parameterize_evaluators | Iterable of evaluators | [ ] | List of evaluation metrics. |
| **Others** | | | |
| **Load()** | | | |
| project_dir | String or path | - | Directory to save project data in. |
| training_project_dir | Path | - | Directory that has the model to be deployed. |
| **Train()** | | | |
| batch_size | Integer | 64 | Number of training samples used through the network during one forward and backward pass. |
| max_epochs | Integer | 32 | Absolute maximum number of epochs to train. |
| lr | Float | 2e-3 | Initial learning rate. |
| num_workers | Integer | 0 | Number of workers to use in the data loaders. Set to 0 on windows! |
| tfms | Callable | field(default_factory=lambda:aug_transforms(flip_vert=True,max_rotate=360, max_zoom=1, max_warp=0, size=224)) | Transforms to apply to the data. |
| metrics | Iterable of callables | field(default_factory=lambda: [BalancedAccuracy()]) | Metrics to calculate on the validation set each epoch. |
| patience | Integer | 3 | Number of epochs without improvement before stopping the training. |
| monitor | String | “valid_loss” | Metric to monitor for early stopping. |
| **multi_input.Train()** | | | |
| batch_size | Integer | 64 | Number of training samples used through the network during one forward and backward pass. |
| max_epochs | Integer | 10 | Absolute maximum number of epochs to train. |
| lr | Float | 2e-3 | Initial learning rate. |
| num_workers | Integer | 0 | Number of workers to use in the data loaders. Set to 0 on windows! |
| tfms | Callable | aug_transforms(flip_vert=True, max_rotate=360, max_zoom=1,max_warp=0, size=224) | Transforms to apply to the data. |
| metrics | Iterable of callable types | [BalancedAccuracy()] | Metrics to calculate on the validation set each epoch. |
| patience | Integer | 3 | Number of epochs without improvement before stopping the training. |
| monitor | String | “valid_loss” | Metric to monitor for early stopping. |
| conts | Iterable of string type | [ ] | List of continuous variables from the clinical table to add to the training. |
| cats | Iterable of string or Category type | [ ] | List of categorical variables from the clinical table to add to the training. |

**Suppl. Table 1: Parameters of different functionalities of DeepMed. * “Map Type” includes e.g. dictionary, key: string, value: map type where key: string and value: any data type.**

| **Metric** | **Options** | **Data type** | **Default** | **Description** |
| --- | --- | --- | --- | --- |
| auroc | not parameterizable | | | Measures the one-vs-rest AUROC for each class of the target label. |
| count | not parameterizable | | | Measures the number of testing instances for each class. |
| p_value | not parameterizable | | | Measures p value for two-tailed t test comparing different classes of the target. |
| r2 | not parameterizable | | | Measures the coefficient of determination. |
| **AggregateStats()** Accumulates stats from subdirectories. | | | | |
|  | label | String | None | Name to be given to the newly aggregated column. |
|  | over | Iterable of string or integer type | None | Index columns to aggregate over. |
|  | conf | Float | 0.95 | Confidence interval calculated during aggregation. |
| **ConfusionMatrix()** Generates a confusion matrix for each class label. | | | | |
|  | min_tpr | Float | None | Minimum true positive rate the confusion matrix shall have for each class. If None, the true positive rate maximizing the F1 score will be calculated. |
| **F1()** F1 Score | | | | |
|  | min_tpr | Float | None | Minimum true positive rate of each class for F1 score. If not given, a threshold which maximizes F1 is used; otherwise, the threshold which yields a tpr of at least min_tpr is used. |
| **Grouped()** Calculates a metric with the data grouped on an attribute. | | | | |
|  | by | String | “PATIENT” | Label to group the predictions by. |
| **Heatmap()** Creates the heatmap of WSI based on tile predictions. | | | | |
|  | wsi_paths | Iterable of path or string type | None | Directory containing the WSIs (needed for superimpose). |
|  | wsi_suffixes | Iterable of string type | [“.svs”, “.ndpi”] | The suffixes of WSIs whose heatmaps to be produced |
|  | superimpose | Boolean | False | Overlay the heatmap onto a low-resolution image of the WSI. |
|  | format | String | “.svg” | The format to save heatmaps. |
| **Roc()** Creates a one-vs-all ROC curve plot for each class. | | | | |
| **TopTiles()** Generates a grid of the best scoring tiles for each class. | | | | |
|  | n_patients | Integer | 4 | Number of best-scoring patients. |
|  | n_tiles | Integer | 4 | Number of best-scoring tiles of each best-scoring patient. |
|  | patient_label | String | “PATIENT” | Column of clinical and slide tables containing patient IDs. |
|  | best_patients | Boolean | True | Whether to select the best or worst n patients. |
|  | best_tiles | Boolean | None | Whether to select the highest or lowest scoring tiles. If set to “None”, then the same as “best_patients”. |
|  | save_images | Boolean | False | Whether to save the tiles separately. |

**Suppl. Table 2: Non-parameterizable and parameterizable evaluation metrics.**
